# *tp53* R217H and R242H Mutant Zebrafish Exhibit Dysfunctional p53 Hallmarks and Recapitulate Li-Fraumeni Syndrome Phenotypes

**DOI:** 10.1101/2024.07.31.605368

**Authors:** Kim Kobar, Lissandra Tuzi, Erin Burnley, Jennifer A. Fiene, Kristianne J. C. Galpin, Craig Midgen, Brianne Laverty, Vallijah Subasri, Timmy T. Wen, Martin Hirst, Michelle Moksa, Annaick Carles, Qi Cao, Adam Shlien, David Malkin, Sergey V. Prykhozhij, Jason N. Berman

**Affiliations:** Children’s Hospital of Eastern Ontario Research Institute, Ottawa, ON, Canada; Department of Cellular and Molecular Medicine, University of Ottawa, Ottawa, ON, Canada; Translational and Molecular Medicine Program, University of Ottawa, Ottawa, ON, Canada; Department of Pathology, IWK Health Centre, Halifax, NS, Canada; Genetics and Genome Biology Program, Hospital for Sick Children, Toronto, ON, Canada; Department of Medical Biophysics, University of Toronto, Toronto, Canada; Peter Munk Cardiac Center, University Health Network, Toronto, ON, Canada; Laboratory of Medicine and Pathobiology, University of Toronto; Department of Microbiology and Immunology, Michael Smith Laboratories, UBC, Vancouver, Canada; Canada’s Michael Smith Genome Sciences Centre, BC Cancer, Vancouver, Canada; Department of Pediatrics, Division of Hematology/Oncology, Hospital for Sick Children, Toronto, ON, Canada; Department of Pediatrics, University of Ottawa, Ottawa, ON, Canada

**Keywords:** Li-Fraumeni syndrome, p53, *tp53*, zebrafish, cancer, cancer predisposition

## Abstract

Li-Fraumeni syndrome (LFS) is a hereditary cancer predisposition syndrome associated with a highly penetrant cancer spectrum characterized by germline *TP53* mutations. We characterized the first LFS zebrafish hotspot mutants, *tp53* R217H and R242H (human R248H and R273H), and found these mutants exhibit partial-to-no activation of p53 target genes, have a defective G1 cell-cycle checkpoint, and are resistant to apoptosis. Spontaneous tumor development histologically resembling human sarcomas developed as early as 6 months. *tp53* R242H mutants had a higher lifetime tumor incidence compared to *tp53* null and R217H mutants, suggesting it is a more aggressive mutation. We observed mutation-specific tumor phenotypes across *tp53* mutants with associated diverse transcriptomic and DNA methylome profiles, impacting metabolism, cell signalling, and biomacromolecule synthesis and degradation. These *tp53* zebrafish mutants demonstrate fidelity to their human counterparts and provide new insights into underlying tumorigenesis mechanisms and kinetics, which will guide targeted therapeutics for LFS.

## Introduction

Li-Fraumeni syndrome (LFS: OMIM 151623) is an autosomal dominant cancer predisposition syndrome with an estimated prevalence of 1:3555 to 1:5476^1^. Tumor onset is typically much earlier than in individuals who have sporadic tumors with somatic *TP53* mutations, often occurring during childhood, and characterized by a diverse tumor spectrum with five “core” cancer types most frequently observed: breast cancer, soft-tissue sarcomas, brain tumors, adrenocortical carcinomas, and bone sarcomas^1,2^. LFS is highly penetrant with a >70% lifetime risk of cancer in males and >90% lifetime risk in females and survivors of their first cancer face an 83-fold higher relative risk of developing subsequent malignancies, likely contributed by treatment of the initial tumor^3^. A clearer understanding of the molecular mechanisms and pathways driving tumor development are critical for the design and delivery of more specific biologically targeted treatments to this highly vulnerable population.

*TP53* is the most frequently mutated gene in cancer and germline *TP53* mutations have been identified in more than 85% of individuals with LFS^1,2^. Over 80% of *TP53* mutations occur as missense mutations in the DNA binding domain, clustering at several “hotspot” residues (175, 245, 248, 273, and 282) in both LFS and sporadic tumors, although at different frequencies^2^. Mutated *TP53* frequently has a dominant-negative effect on wildtype p53 and can also acquire novel functions in the form of gain-of-function (GOF) mutations^4^. p53 is known as the “guardian of the genome” for the critical role it serves as a tumor suppressor. Following a cellular stress event, like DNA damage, wildtype p53 rapidly accumulates in the cell and activates numerous target genes to regulate key cellular protective processes, such as initiating DNA repair, inducing cell cycle arrest, and promoting apoptosis^5^. Additionally, wildtype p53 exerts tumor-suppressive functions by preventing major epigenetic alterations in order to maintain appropriate DNA methylation levels, and loss of function *TP53* mutations are associated with hypomethylation across multiple cancer types^6–10^. Thus, further evaluating how *TP53* mutations affect the epigenetic landscape could identify novel therapeutic and diagnostic strategies for LFS and sporadic cancers.

In this study, we present the phenotypes of the first two zebrafish *tp53* hotspot mutants, R217H and R242H (human R248 and R273H hotspot residues, respectively), generated in the transparent *casper* background using a CRISPR/Cas9-based approach to knock in the desired point mutations^11,12^. We found that these mutants have lost several p53 tumor suppressive functions as they have decreased expression of p53 target genes; loss of wildtype p53 functionality as they develop normally following *mdm2* knockdown; are resistant to p53-mediated apoptosis; have a malfunctioning G_1_ cell-cycle checkpoint; and mutant proteins show dominant negative activity. These mutant alleles result in tumor development beginning at 6 months post-fertilization (mpf) that histologically resemble human sarcomas with heterozygote tumors displaying loss of *tp53* heterozygosity (LOH). We observed mutant-specific differences with regard to tumor onset, anatomic location, lifetime incidence, and sex bias, collectively suggesting GOF activity in the point mutants as compared to the *tp53* null line^13^. Both RNA and whole genome bisulfite sequencing (WGBS) on *tp53* mutant larvae revealed transcriptomic and DNA methylome changes that impacted many cellular processes including metabolism, cell signaling, and biological macromolecule synthesis, degradation, and transport.

## Results

### Generation and validation of tp53 R217H and R242H CRISPR Knock-In Mutants

The human p53 R248 and R273 hotspots (zebrafish R217 and R242, respectively) are the two most commonly mutated germline and somatic p53 residues in cancer^2^. Located in the DNA binding domain (Supplementary Figure S1A), these mutations are referred to as contact mutations as they interfere with the ability of p53 to bind DNA^14^. As these specific residues and the areas around them are highly conserved (Supplementary Figure S1B), we generated zebrafish *tp53* R217H/R217H and *tp53* R242H/R242H (hereafter referred to as R217H and R242H unless specifically mentioned as heterozygous) using a CRISPR ssODN knock-in approach as previously described by our group^11^ (Supplementary Figure S1C).

To assess both the stability and quantity of the mutant p53 proteins, we dosed the embryos with 30 Gray (Gy) of irradiation (IR) as p53 protein levels are typically very low in the absence of a cellular stress event. We performed western blots on embryos 6 hours post-IR (hpi) (30 hpf) and observed that *tp53* null zebrafish completely lacked p53 protein, while both R217H and R242H mutant protein levels were increased compared to *casper* for both normal and IR conditions (Supplementary Figure S1D,E).

### tp53 mutants have decreased expression of p53 target genes

A cellular stress event, like DNA damage, leads to the accumulation of the p53 protein which binds to and activates numerous target genes to initiate tumor suppressive functions^15^. To induce the p53 pathway, we treated 24 hpf embryos with either 30 Gy using IR or 0.1 μM Camptothecin (CPT), a topoisomerase I inhibitor that causes DNA damage, employing 0 Gy and DMSO treatment as controls, respectively. To investigate if the activation of these target genes was perturbed in the *tp53* mutants, we performed qPCR on *casper* and *tp53* mutant embryos to measure the expression of several p53 target genes: cell cycle genes: *ccng1* (cyclinG) and *cdkn1a* (p21); apoptosis promoting genes: *bbc3* (puma) and *mir34a*; and p53 regulatory genes: *mdm2* and *tp53* (Figure 1, Supplementary Figure S2).

**Figure 1.**
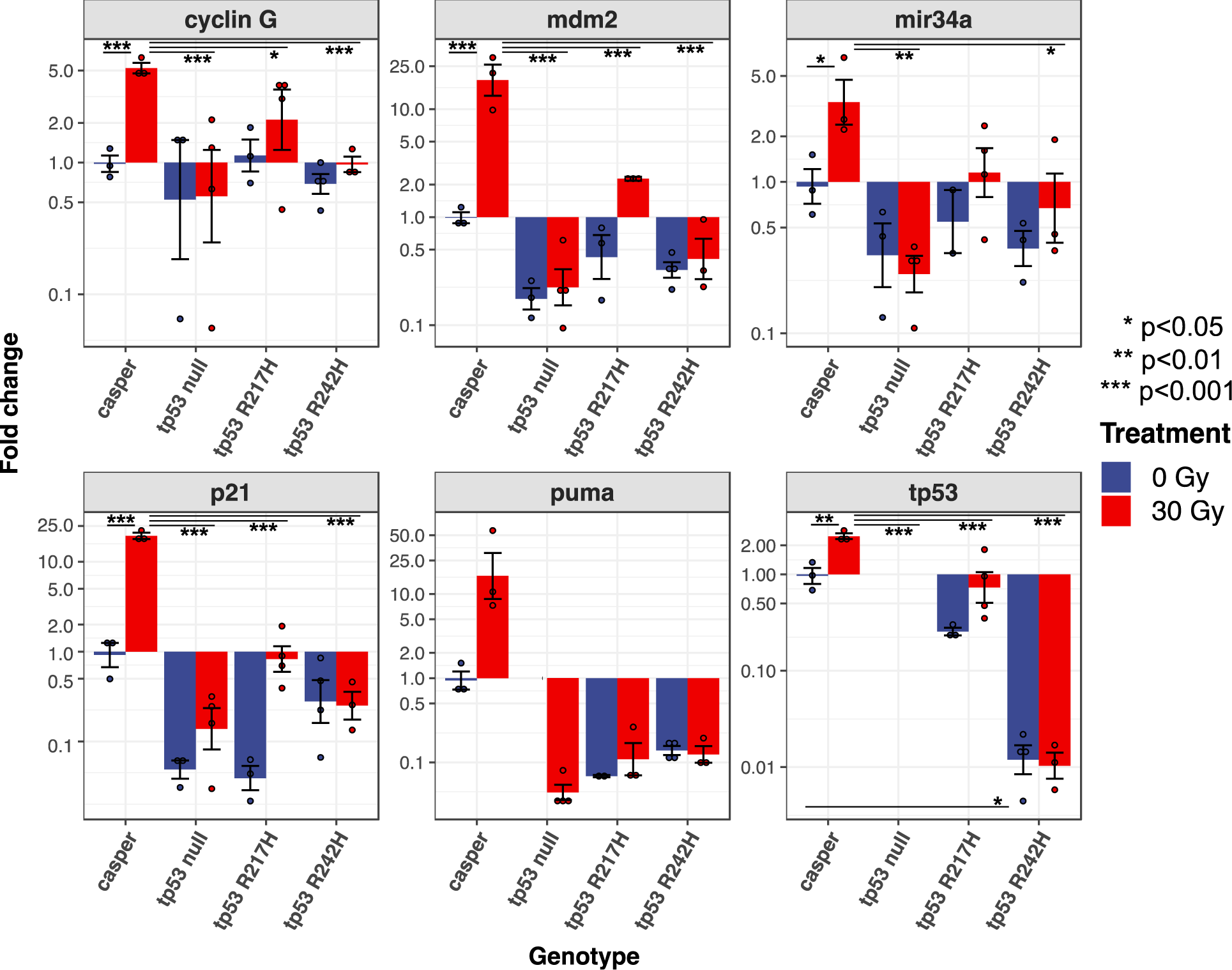
The fold change of expression of several p53 target genes following induction of the DNA damage pathway is decreased in *tp53* mutants. Thirty 24 hpf embryos per sample were exposed to 0 or 30 Gy of IR and were collected at 30 hpf (6 hpi). Expression was not detected in the *tp53* null 0 Gy *puma* and 0/30 Gy *tp53* groups. Data represents 2-4 technical replicates and 3-4 biological replicates. The fold change was normalized to the *casper* 0 Gy group for each gene. Outliers were identified and excluded using Grubbs’ test with alpha <0.05. Statistical significance was determined by a two-way ANOVA and a Tukey’s Honest Significant Difference test. Error bars represent standard deviation. The Y-axis is logarithmic scale 10.

All six p53 target genes were strongly upregulated in both the *casper* CPT and 30 Gy samples, while all three *tp53* mutants had decreased expression of each target gene compared to *casper* upon p53 pathway induction (Supplementary Figure S2, Figure 1). The *tp53* null and R242H mutants displayed similarly decreased expression in all genes with almost no activation of p53 target genes upon CPT (two-way ANOVA p-value <0.001 for all six genes) or IR treatment (two-way ANOVA p-value <0.001 for *ccng1, mdm2, p21,* and *tp53*), indicating that the R242H mutation interferes with the ability of p53 to induce transcription of these target genes. By contrast, the CPT-treated R217H mutants had partial activation of all six genes compared to the CPT-treated *casper* groups, suggesting it may be a hypomorphic mutation with a less severe phenotype (Supplementary Figure S2). Interestingly, this trend was not as strong in the IR R217H mutants; however, *puma* was the only gene for which IR had no impact (Figure 1). Notably, all genes except *cyclinG* had much lower baseline expression levels in the R217H group compared to *casper.* Altogether, this data suggests that R217H mutants may be more sensitive to CPT treatment, or a p53-independent CPT-induced mechanism may be present. Importantly, *tp53* null mutants had no *tp53* expression across all treatment groups as well as no detectable *puma* expression in the 0 Gy group (Figure 1, Supplementary Figure S2).

### tp53 R217H and R242H mutants develop normally following mdm2 knockdown

MDM2 is an important regulator of the p53 pathway as it is both a p53 target and an E3 ubiquitin ligase that binds to and promotes the degradation of p53 in a negative feedback loop^16^. The complete loss of Mdm2 in mice is embryonically lethal as it results in over-accumulation of p53; however, these mice survive if they also lack p53^17–19^. This was similarly observed in zebrafish using a splice-blocking morpholino to knockdown *mdm2* mRNA accumulation which increased apoptosis, cell cycle arrest, and lethality, and these phenotypes were not observed upon simultaneous knockdown of *tp53* and *mdm2*^20^. Thus, inhibiting *mdm2* serves as a direct test of whether the mutant p53 protein retains any key functions of the wildtype p53, such as induction of apoptosis and cell cycle arrest^20–22^.

To functionally assess if the zebrafish mutant R217H or R242H proteins retained any wildtype function, we knocked down *mdm2* using a previously well-characterized morpholino ^20^ and assessed the resulting phenotypes at 72 hpf (Figure 2). We observed the presence of *mdm2* knockdown-associated phenotypes, such as high levels of deformities, development delay, and death, only in the *casper* injected group (Figure 2). As expected, the MO-injected *tp53* null mutants developed normally, as they completely lack the p53 protein. Additionally, MO-injected R217H and R242H embryos developed normally, indicating that R217H and R242H mutant p53 proteins have lost wildtype functionality, as partial p53 functionality would likely still cause lethality. A small subset of both control and MO-injected *tp53* mutant embryos showed abnormal development, indicated as “not representative” in the ratios, however, these were present in both the control and injected groups and were not nearly as severe as the phenotypes observed in the MO-injected *casper* embryos (Figure 2). Incorrect *mdm2* splicing was verified by two independent PCRs on pooled *mdm2* MO-injected embryos (Supplementary Figure S3).

**Figure 2.**
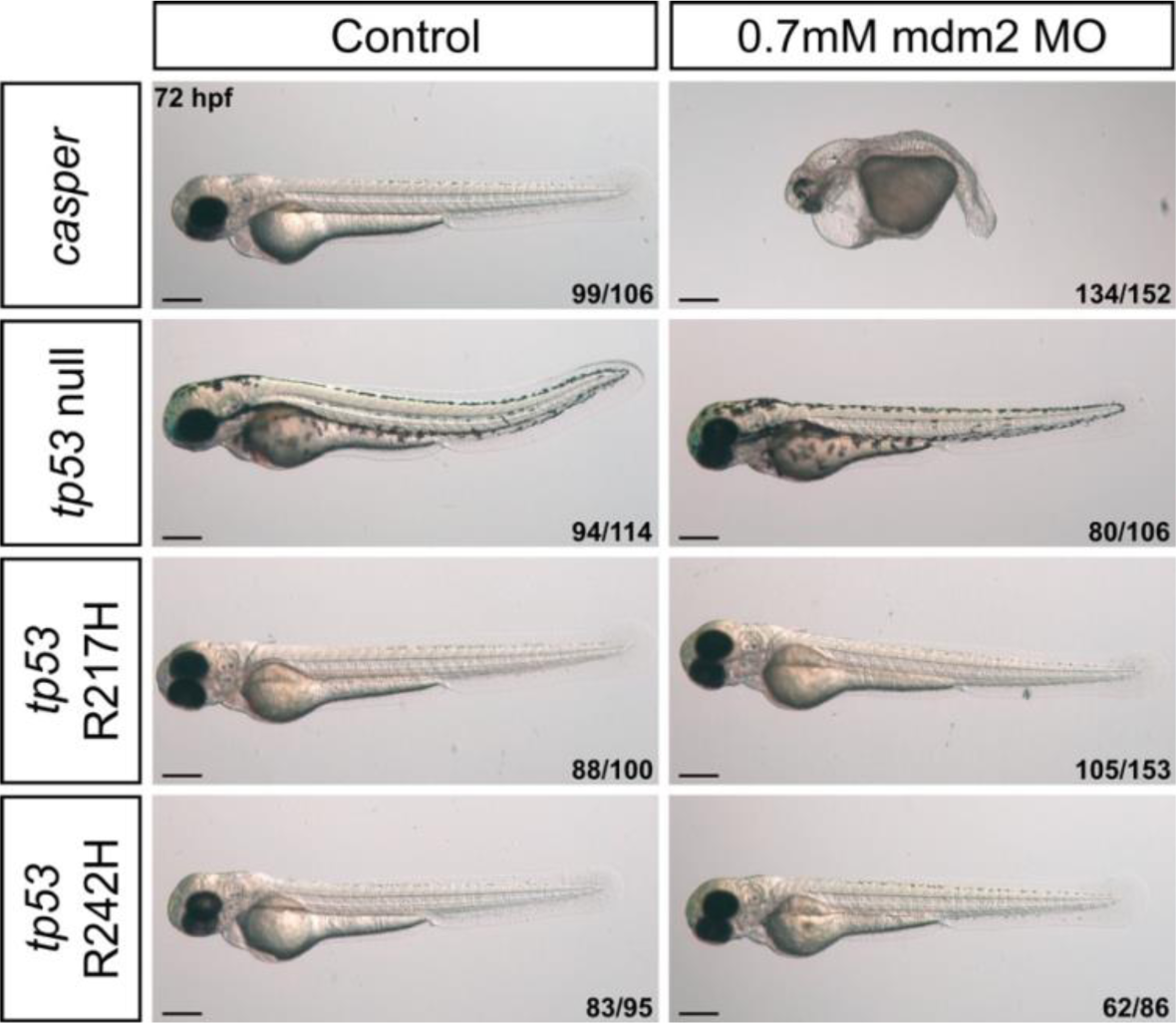
mdm2 MO-injected mutant *tp53* embryos are resistant to mdm2 loss-associated phenotypes due to mutant p53 LOF. Single cell embryos were injected with 0.7mM of a splice-blocking *mdm2* morpholino. Images were captured at 72hpf. Representative images shown for each group as described by ratio in bottom right corner of each image, from four independent experiments. Scale bars, 300μM.

### tp53 mutants are resistant to p53-mediated apoptosis in a dominant-negative manner

Next, we investigated the ability of R217H and R242H mutants to elicit an apoptotic response. We performed acridine orange (AO) staining, which is a stain used in live animals that identifies apoptotic cells in zebrafish typically in the neural tube along the dorsal side of the embryo^20,23–25^. An apoptotic pattern was only observed in the *casper* 30 Gy group, revealing that all three p53 mutants were resistant to p53-mediated apoptosis (Figure 3A).

**Figure 3.**
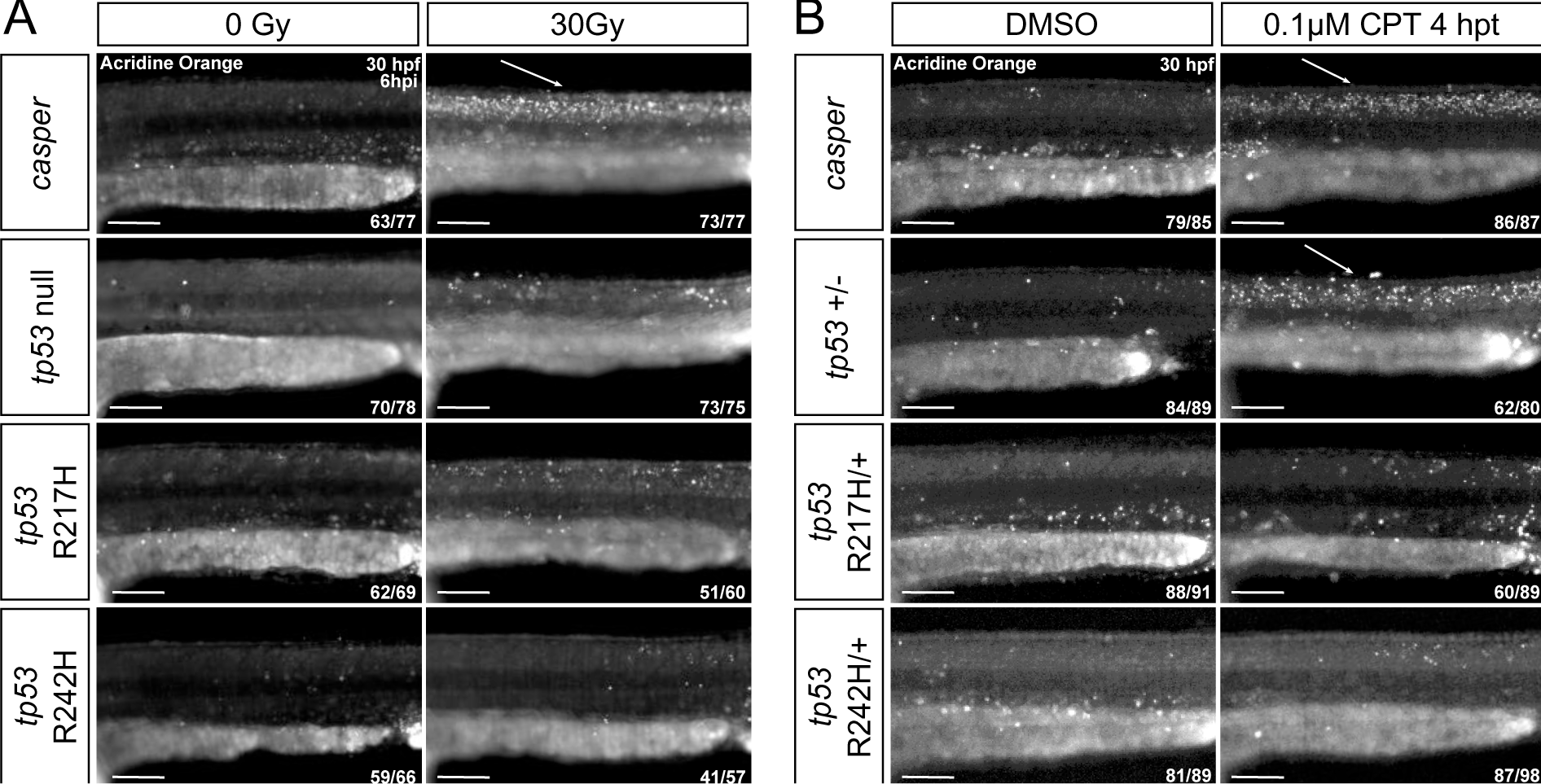
*tp53* zebrafish mutant embryos are resistant to apoptosis and exhibit dominant negative activity following induction of the DNA damage pathway. (A) Embryos were exposed to +/- 30Gy of IR at 24 hpf and stained with AO at 6 hpi. (B) 26 hpf embryos were treated with 0.1μM CPT or DMSO for 4 hours and stained with AO at 30 hpf. The white arrows highlight the typical apoptotic pattern observed along the dorsal spine. Images are of the dorsal spine above the yolk extension. Representative images shown for each group as described by the ratio in bottom right corner of each image. n=3 replicates. Scale bars, 100μM.

Several of the most common *TP53* missense mutant proteins have been identified to have dominant-negative activity, including the human R248 and R273 mutants, where the mutant protein blocks the functional activity of the wildtype protein if present^26,27^. To investigate this phenomenon in the zebrafish context, we performed AO staining on 30 hpf *casper* and heterozygous *tp53* mutant embryos treated with 0.1 μM CPT for 4 hours to induce the p53 pathway (Figure 3B). An apoptotic pattern was observed in the *casper* and heterozygous *tp53* null treated embryos, as the heterozygous *tp53* null embryos possess one copy of wildtype p53 which was sufficient to induce apoptosis (Figure 3B). However, heterozygous R217H and R242H mutants displayed a p53-induced apoptosis-resistant phenotype reminiscent of their homozygous counterparts, indicative of a dominant negative effect for both mutations (Figure 3B).

### tp53 mutants have a defective G_1_ cell cycle checkpoint

Inducing cell cycle arrest and inhibiting proliferation upon a DNA damage event is another critical role of the p53 pathway^28^. To evaluate these functions in the *tp53* mutants, we performed whole-mount immunofluorescence staining for phospho-S10 histone H3 (pH3), a proliferative marker of cells in the G_2_/M phase, on 28 hpf (3 hpi) embryos that were exposed to 0 or 30 Gy at 25 hpf (Figure 4A). While all genotypes experienced a reduction of proliferating cells following IR, all three *tp53* mutants demonstrated a modestly greater number of proliferating cells following 30 Gy exposure compared to *casper* controls (Figure 4A).

**Figure 4.**
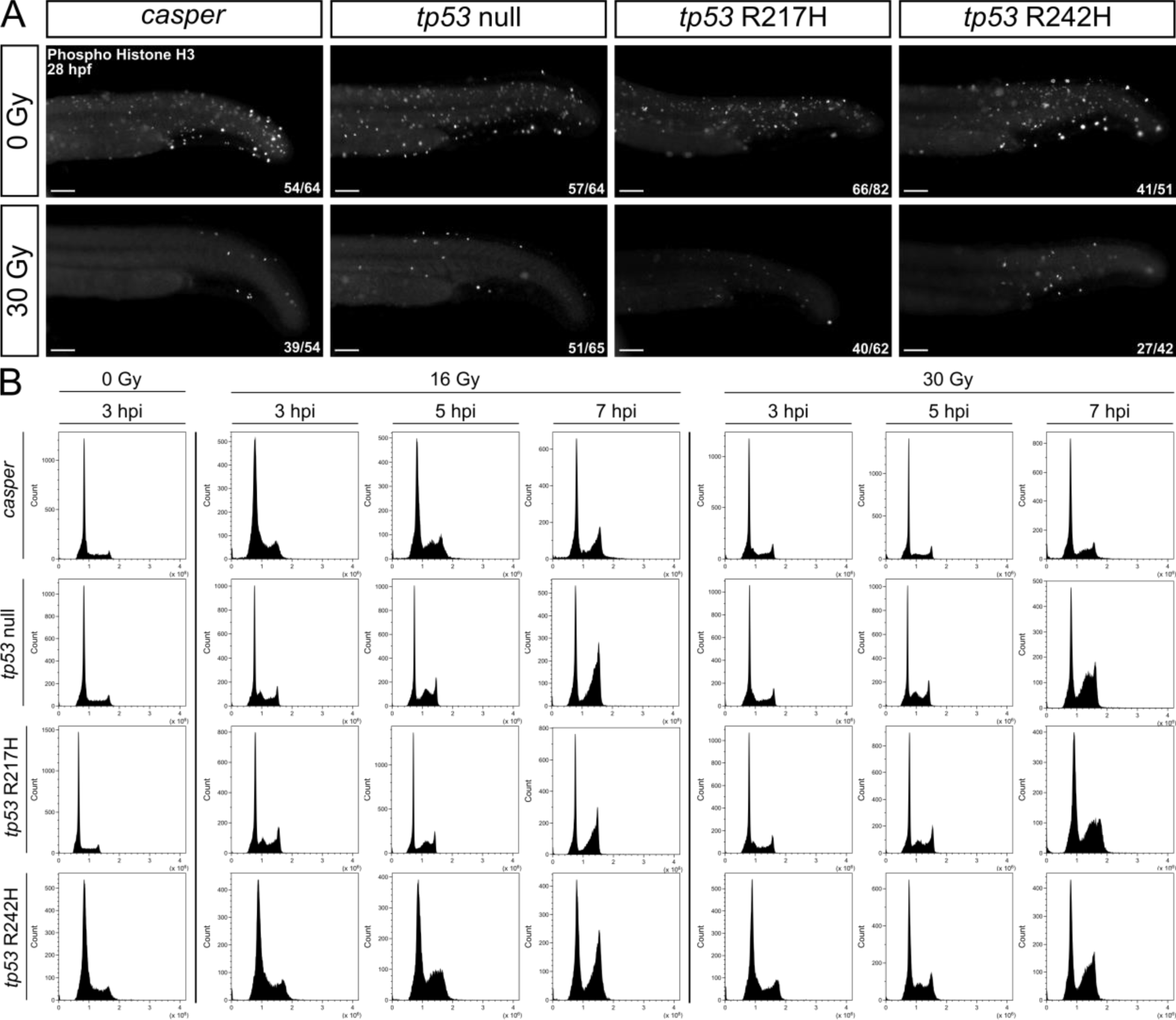
*tp53* mutant embryos display increased proliferating cells and a defective G_1_ cell cycle checkpoint following IR compared to *casper* embryos. (A)) Immunofluorescence for PH3 performed on 28hpf (3hpi) embryos +/- 30Gy. Representative images shown for each group as described by ratio in bottom right corner of each image. n=3 replicates. (B) Flow cytometry histograms of DNA content in each cell cycle phase (G_1_, S, G_2_/M) at 3, 5, and 7 hpi dosed with 0, 16 or 30 Gy in *casper, tp53* null, R217H, and R242H embryos. Embryos were enzymatically dissociated, fixed, permeabilized, and stained with FxCycle violet. Each plot was gated to 25000 cells (+/-100 cells). Results are representative of three independent experiments. Scale bars, 100μM.

To investigate the functionality of the cell cycle checkpoints in the *tp53* mutants, we performed flow cytometry measuring DNA content on embryos dosed at 24 hpf with 0, 16, or 30 Gy and collected at 3, 5, and 7 hpi (Figure 4B). For all genotypes in the 16Gy treatment group, there was an accumulation of cells arrested in the G_1_/early S phase at 3 hpi that moved into the mid-S phase at 5 hpi and arrested in the G_2_/M phase at 7hpi (Figure 4B). Strikingly, *tp53* null, R217H, and R242H embryos in the 30 Gy treatment groups exhibited an accumulation of cells in mid-S phase at 5 hpi and in G_2_/M at 7 hpi, while the *casper* embryos retained most of their cells in the G_1_ phase. This signifies that the p53-dependent G_1_ checkpoint is defective in the *tp53* mutants, while the G_2_/M checkpoint remains intact.

### tp53 mutants spontaneously develop tumors with mutation-specific phenotypes

Altogether, the *tp53* mutants lack many wildtype p53 tumor suppressive functions which predisposed them to develop spontaneous tumors as adults beginning at 6 mpf (Figure 5). We conducted a long-term tumor evaluation study until 16 mpf and observed tumor development in multiple locations throughout the fish (Figure 5). H&E staining on a subset of tumors (n=15) revealed they were all sarcomas, one of the predominant LFS tumors, with two distinct morphologies reminiscent of either small-round-blue-cell (SRBC) or spindle cell-like tumors, and three highly vascularized tumors (Figure 5A, Supplementary Table S1). Several of the tumors with spindle cell-like histology were consistent with zebrafish malignant peripheral nerve sheath tumors (zMPNSTs)^13,20,29^. Many tumors had variations of these morphologies, such as the presence of rosettes, a more diffuse spindle cell-like appearance, or even features of both SRBC and spindle cell-like patterns within the same tumor (Figure 5A, Supplementary Table S1). Remarkably, one R217H flank tumor had unique histologically which resembled human alveolar rhabdomyosarcoma (aRMS) (Supplementary Table S1). We were limited with further classification of these tumors due to a lack of cross-reacting human protein-directed antibodies, preventing immunohistochemical evaluation in these zebrafish^20^.

**Figure 5.**
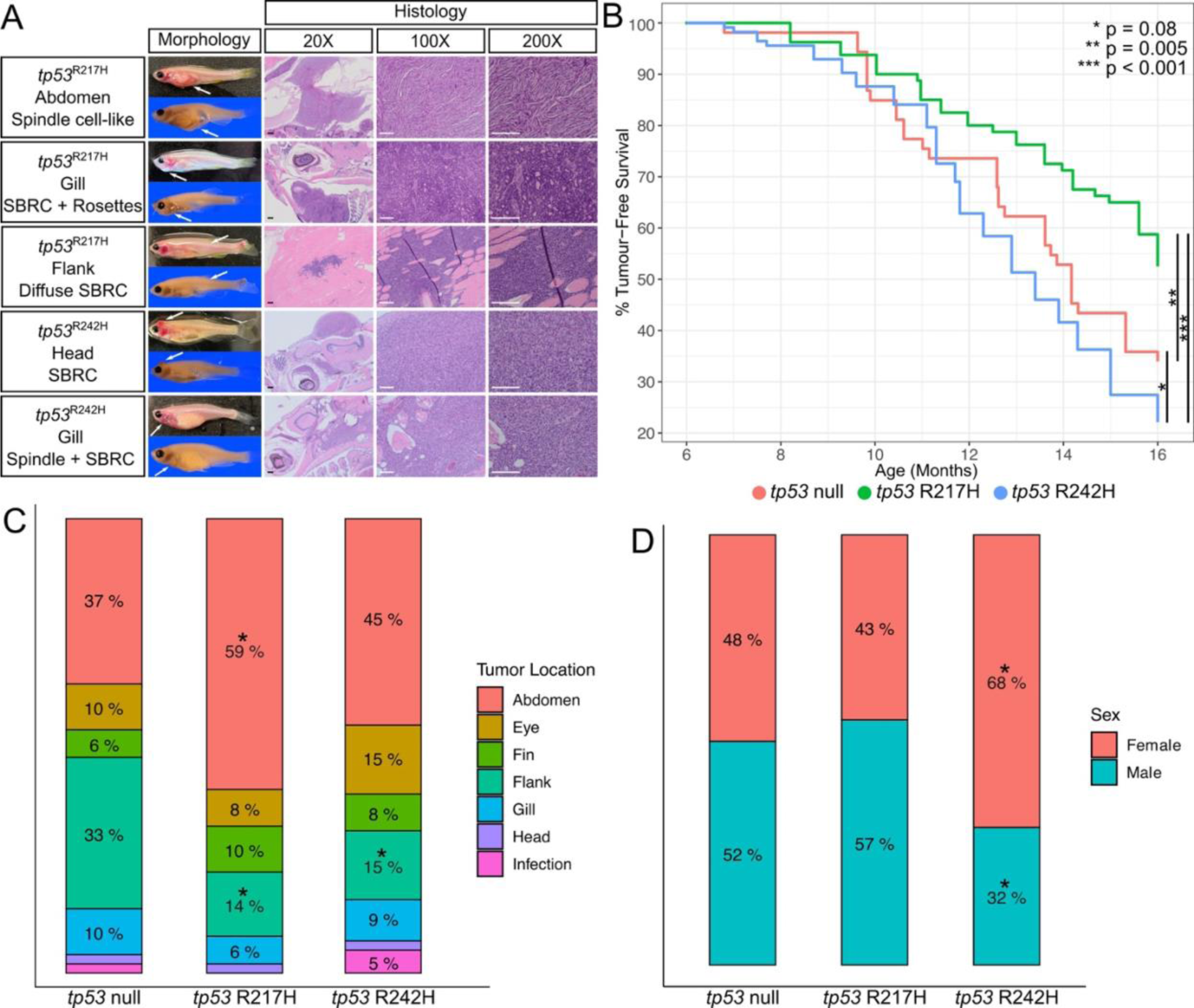
LFS zebrafish mutants develop sarcoma-like tumors with a divergent anatomical distribution of tumor development and higher lifetime tumor incidence in the *tp53* R242H mutants. (A) Live and fixed morphology of *tp53* mutant tumours, indicated by white arrows, and histology panels represent H&E stained sections of the anatomical area of the tumor. (B) Kaplan-Meier analysis showing tumor-free survival and onset in the *tp53* null (n=52), R217H (n=77), and R242H (n=112). Stars show significance as *tp53* null vs. R217H – p=0.005, *tp53* null vs. R242H – p=0.08, R217H vs. R242H – p = 1×10^-7^. Percent tumor-free survival refers to the age a specific fish had died or was euthanized due to tumor development. Tumor development was not observed in the *casper* line during this time period. Scale bars for 20X are 200μM. Scale bars for 100X and 200X are 100μM. (C) Comparison of anatomical site of tumor development in *tp53* null (n=51), R217H (n=49), and R242H (n=117) that developed tumors. The infection category represents fish with severe infection phenotypes (dropsy and edema) and may indicate the presence of a leukemia/lymphoma. A “*” above the percentage indicates a significant difference compared to *tp53* null (p <0.05 – two-tailed two proportions Z-test). (D) Comparison of tumor fish sex across the *tp53* null (n=52), R217H (n=46), and R242H (n=111) mutants. A “*” above the percentage indicates a significant difference compared to *tp53* null (p <0.05 – two-tailed two proportions Z-test).

Additionally, we found that R242H mutants had an earlier time to tumor onset (6-8 mpf, except for one *tp53* null fish which also developed a tumor this early) and a significantly higher lifetime incidence (20% survival at 16 mpf) compared with *tp53* null and R217H mutants (Figure 5B). As R242H mutants experience more aggressive tumorigenesis compared to zebrafish that lack the p53 protein, this suggests that the R242H mutant protein has gained novel pro-tumorigenic functions. By contrast, R217H mutants had a lower lifetime tumor incidence compared to the other *tp53* mutants (Figure 5B), which combined with the p53 target gene expression data (Figure 1, Supplementary Figure S2), suggests a partial retention of wildtype p53 functions delaying tumor onset.

Tumors arose in many anatomic locations across all three *tp53* mutants, including the abdomen, eyes, fins, flank, gills, and head (Figure 5C, Supplementary Figure S4). The head and certain flank tumors are likely angiosarcomas based on similarities in physical appearance to documented angiosarcomas in *tp53* null mutants^13^. For all mutants, abdominal tumors were the most common, with a significantly higher distribution in the R217H mutants (Figure 5C). Notably, flank tumors were significantly lower in R217H and R242H mutants compared to *tp53* nulls (Figure 5C). Several fish did not exhibit a distinct mass but rather presented with strong indicators of infection, including severe edema, abdominal swelling, and protruding scales with a pinecone-like appearance (Supplementary Figure S4C), which may indicate leukemia/lymphoma, though further analysis was not undertaken. Similarly, *tp53* null fish showing these infection phenotypes were diagnosed using RNA-seq to have an aggressive NK cell-like leukemia^13^. Further, we observed skewed sex ratios in the fish that developed tumors as significantly more R242H females were affected (68% female) compared to *tp53* null (48% female) and R217H (43% female) sex distributions (Figure 5D). Altogether, these observations suggest distinct GOFs associated with the R217H and R242H mutant proteins promoting tumor development as compared to *tp53* null mutant phenotypes.

### tp53 R217H/+ and R242H/+ tumors exhibit loss of heterozygosity

Loss of heterozygosity (LOH) of the wildtype *TP53* allele, often with gain of the mutant allele, is frequently observed in LFS tumors^2,30^. To investigate if the heterozygous *tp53* mutants exhibit LOH, we compared tumor DNA with a paired fin-clip to provide genomic DNA. We found LOH in 4/5 *tp53* R217H tumors and 4/4 *tp53* R242H tumors that were tested, as the fin-clip DNA was heterozygous while the tumor DNA was homozygous (Supplementary Figure S5B). Contamination of non-tumor tissue taken with the tumor tissue sample may have limited the LOH diagnosis for the one negative tumor. Tumors first arose in *tp53* R217H/+ mutants at ∼7 mpf compared to ∼8 mpf in *tp53* R242H/+ fish, with *tp53* R217H/+ having a higher lifetime incidence at ∼40% compared to ∼20% in *tp53* R242H/+ mutants (Supplementary Figure S5C).

### Altered gene expression in the tp53 mutants leads to changes in metabolism, biosynthetic and developmental processes, and cell signalling

Key genes and pathways in *tp53* zebrafish mutants that are differentially expressed may contribute to the mutation-specific heterogeneity observed in tumor phenotypes. We performed bulk RNA sequencing on pooled 30 hpf embryos of each genetic background (*casper*, *tp53* null, R217H, or R242H) exposed to either 0 or 30 Gy (6 hpi). This time point will enable the identification of dysregulated developmental mechanisms that may predispose to tumor development, as well to investigate the transcriptional consequences following p53 pathway induction. From these samples, we removed one *tp53* null 30 Gy sample and one *tp53* R217H 0 Gy sample as technical outliers as these samples each had very high expression of subsets of genes with overall differing expression from their biological replicates (Supplementary Figure S6).

To determine how the p53 signaling pathway was impacted in our mutants, we mapped predicted p53 target genes^31^ to their zebrafish orthologues, representing 400 genes, and found 19 of these were differently expressed between the 0 and 30 Gy treatment groups across each genotype (Figure 6A). There is a clear response following IR in the *casper* group that is not present for *tp53* null, R217H, or R242H, indicating that the p53 signaling pathway is not functional in totality in these mutants (Figure 6A). The target genes validated with qPCR (Figure 1 - indicated in red – *ccng1, bbc3, mdm2, cdkn1a,* and *tp53*) exhibited a similar expression pattern (Figure 6A).

**Figure 6.**
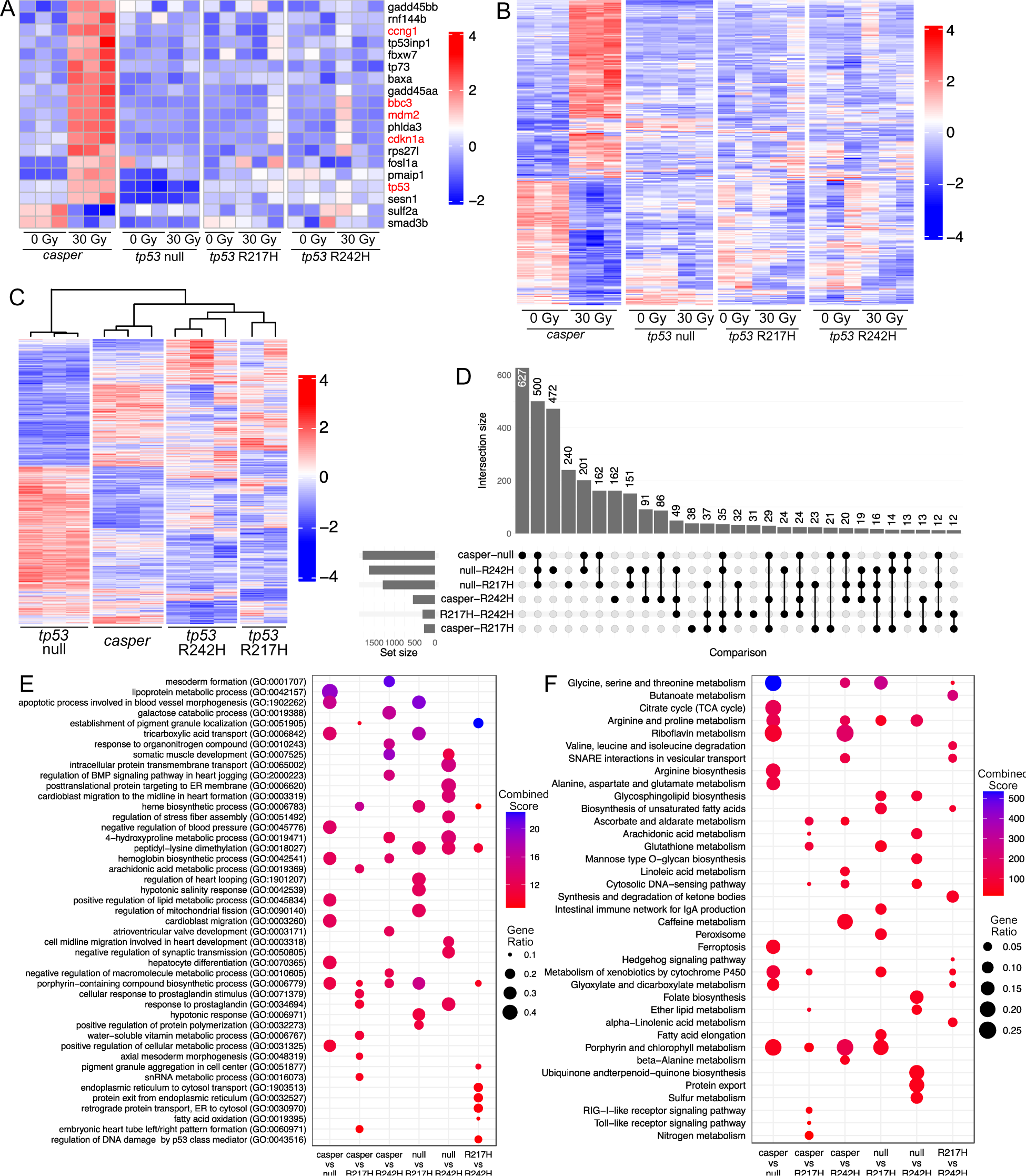
IR treatment impacts gene expression in *casper* but not *tp53* mutant embryos while under control conditions, *casper, tp53* null, R217H, and R242H mutants have largely varying DEGs and GO terms affecting metabolism, biosynthesis and developmental processes, and cell signalling. (A) Heatmap of p53 target genes that were differentially expressed between 0 and 30 Gy conditions across all genotype/treatment samples (adjusted p-value <0.05, fold change >2). Genes that were also measured by qPCR in Figure 2 are listed in red. (B) A heatmap showing DEGs from all genotype comparisons from the 0 Gy treatment groups (adjusted p-value <0.05, fold change >2). (C) A heatmap of all differentially expressed genes from all genotype/treatment group comparisons (adjusted p-value <0.05, fold change >2). (D) UpSet plot shows the number and overlap of DEGs between each genotype from the 0 Gy treatment groups. The total number of DEGs per set is shown on the x-axis and the y-axis shows the intersection of DEGs for each comparison. Comparisons with an intersection of less than ten are not shown. (E, F) Dot plots showing the top ten GO BP (E) and KEGG pathway terms (F), ranked by combined score, for each comparison under control conditions. Dot size corresponds to the ratio of DEGs per total number of genes for each term and the colour corresponds to the combined score.

Next, we looked at all differentially expressed genes (DEGs) between the 0 and 30 Gy groups for each genotype to determine the impact of IR on gene expression. As expected, IR led to vast changes in *caspers* with 273 DEGs while there were only 6, 7, and 14 DEGs between *tp53* null, R217H, and R242H treatment groups, respectively (Figure 6B, Supplementary Figure S7A). The only similar DEG across all genotypes was *cdca7a* (cell division cycle associated 7), a c-Myc-responsive gene associated with tumorigenesis^32^. Additionally, we applied GO enrichment analyses to understand the impact of IR treatment on KEGG pathways (Supplementary Figure S7B) and Biological Processes (BP) (Supplementary Figure S7C) for each genotype. As expected, IR treatment in *caspers* led to changes in the p53 pathway response, apoptosis, and DNA replication and repair with only a minor response in DNA replication and cell cycle in *tp53* null and R242H mutants (Supplementary Figure S7B, C). Altogether, this suggests that p53 pathway functions are conserved in zebrafish and IR treatment does not impact transcription in the *tp53* mutants either globally or in a p53-specific manner.

As we found that IR exposure did not have a significant impact on gene expression in *tp53* mutants, we next examined DEGs under basal, non-stress conditions to investigate mutation-specific genetic underpinnings (Figure 6 C, D). Notably, a 6-set comparison between the 0 Gy *casper*, *tp53* null, R217H, and R242H groups revealed the *tp53* null group was strikingly different with over 2300 DEGs when compared to the other three genotypes; however, there were many diverse groups of DEGs across all genotype comparisons (Figure 6 C, D). GO BP and KEGG pathway analyses identified enriched terms associated with metabolism, notably carbohydrates, fatty acids, amino acids, and their precursors; biological macromolecule synthesis, degradation, and transport of carbohydrates, porphyrin-containing compounds, tricarboxylic/fatty acids, intracellular proteins, and hemoglobin; and cell signaling (Figure 6E, F). Altogether, these altered processes in the *tp53* mutants suggest wide-scale metabolic reprogramming and deregulation of cellular energetics to provide energy and increased bioavailability of substrates needed for the transformation and rapid proliferation of cancer cells.

### tp53 mutants have altered DNA methylation which affects cell signalling, metabolic processes, biomacromolecule synthesis/degradation

p53 plays a key role of maintaining epigenetic stability and *TP53* mutations are strongly associated with aberrant methylation found in cancer. We performed WGBS on 5 dpf *casper, tp53* null, R217H, and R242H larvae under steady state conditions (Figure 7). Similar to RNA sequencing results, we found DNA methylation in the *tp53* null group was very different from *caspers* and *tp53* point mutants (Figure 7A, B). Interestingly, *casper* and *tp53* R242H mutants were the most similar (Figure 7A, B). To identify differently methylated regions (DMR), we applied a false discovery rate <0.05 and >25% methylation difference to 1000 bp windows of DNA and found 150 DMRs across 226 genes as different across at least two genotypes (Figure 7C). GO pathway analysis for the top ten enriched GO BP (Figure 7D) and KEGG pathway (Figure 7E) terms, ranked by combined score, revealed changes in cell signaling and transporter activities, metabolism, and the synthesis and degradation of biological macromolecules. Combined with the RNA sequencing results, the overlapping key themes suggest that cell signalling, metabolism, and the synthesis/degradation of biological macromolecules are vastly altered in the *tp53* mutants and may represent novel GOFs promoting tumorigenesis that could serve as previously unrecognized therapeutic targets.

**Figure 7.**
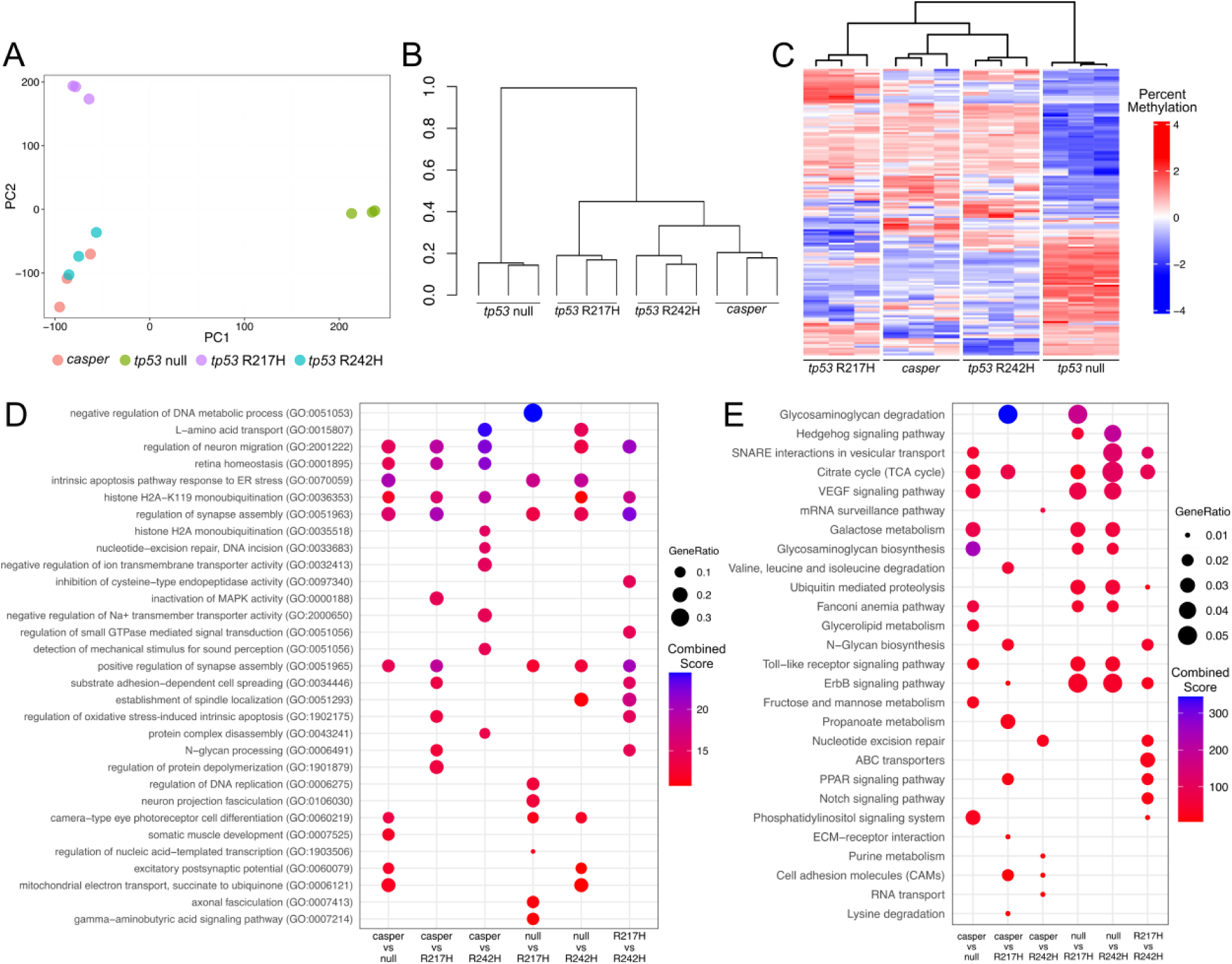
DNA methylation changes in *tp53* mutant zebrafish affects cell signalling, metabolic processes, and biological macromolecule synthesis and degradation. A) Primary component analysis (PCA) plot shows clustering of the bulk WGBS *casper, tp53* null, R217H and R242H pooled 5 dpf zebrafish samples. B) Dichotomous diagram of the WGBS samples using the Ward clustering method with a correlation distance method. C) Heatmap showing the percent methylation of 150 DMRs identified from 1000bp windows with an FDR <0.05 and a methylation differences of >25%. D) Dot plots showing the top ten GO BP (D) and KEGG pathway (E) terms, ranked by combined score, between each genotype comparison. Enrichment analysis was performed on 226 genes that were associated with the 150 DMRs. Dot size corresponds to the ratio of associated genes per total number of genes for each term and the colour corresponds to the combined score.

## Discussion

### R217H and R242H alleles lack wildtype p53 tumor suppressive functions

Altogether, we have shown that the *tp53* R217H and R242H mutants have lost wildtype p53 functions which predisposed spontaneous tumor development in early adulthood. Like wildtype p53, both R217H and R242H mutant protein levels are increased by IR, which indicates that the loss of these tumor suppressive phenotypes is not the result of altered p53 protein stabilization. Interestingly, the mutant protein levels are higher than wildtype under each control and IR conditions. This is a phenomenon often seen in human tumors and was also observed in several mouse models harboring *Tp53* missense hotspot mutations^33,34^.

One of the main ways that p53 elicits its tumor suppressive functions is through the activation of a few hundred key target genes, a process which we showed is disrupted in the *tp53* mutants evidenced by partial-to-no expression of p53 target genes following p53 pathway induction. Additionally, we showed that IR treatment does not significantly influence gene expression on a global level in the *tp53* mutants. As R217H and R242H mutations are both contact mutations that interfere with the ability of p53 to bind to DNA, it is likely that this is the main mechanism by which the mutant proteins have lost the ability to promote wildtype p53 functions^14^. However, R217H mutants retained partial activation of several key p53 target genes which does not appear to have a phenotypic impact following *mdm2* knockdown nor resistance to apoptosis, but over time this partial activation may confer some tumor-suppressive effects that contribute to the less aggressive tumor phenotypes.

Although the *tp53* mutants exhibited minimal-to-no *p21* activation upon IR, suggesting they would not be able to induce cell cycle arrest, they experienced a reduction of proliferating cells similar to *casper* following IR. Interestingly, *cdca7a*, a direct MYC target gene, the downregulation of which is associated with decreased cellular proliferation, was downregulated upon IR treatment in all four genotypes^32^. Notably, *cdca7a* was the only common treatment-associated DEG between *casper*, *tp53* null, R217H, and R242H mutants and may explain this observed p53-independent decrease in proliferating cells following IR in all groups. Of note, *tp53* M214K zebrafish, a loss-of-function (LOF) mutation, showed that the pH3 staining patterns of IR-treated embryos did not differ from those in wildtype embryos, however, M214K embryos also exhibited a defective G_1_ checkpoint following IR^29^.

### tp53 R217H and R242H mutants develop tumors representative of LFS with mutation-specific characteristics

LFS mouse models, including *Tp53* +/-, *Tp53* -/- and several missense “hotspot” mutants develop tumors with differential onset, type, and metastatic potential, with the missense mutants displaying a broader and highly malignant tumor spectra similar to what is observed in LFS patients^1,33–39^. For example, *Tp*53 R270H/+ (human R273H) mice have an increased tumor burden with distinct differences in tumor type including carcinomas and hemangiomas, as well as an increased metastatic potential compared to *Tp53* +/- mutants^34^.

Several zebrafish *tp53* mutants have been generated to investigate cancer predisposition with many lines exhibiting resistance to p53-mediated apoptosis at physiological temperatures and subsequently developing tumors that are histologically similar to their human counterparts^13,20,40,41^. *tp53* M214K (human M246K) zebrafish predominantly developed zMPNSTs in the abdomen and eye with a 28% incidence over 16.5 months with one incidence of melanoma^29^. *tp53* I166T (human I195T) zebrafish developed spindle cell sarcomas in the abdomen, eye, flank, gill, rectum, and skin beginning at 9 mpf with a 50% incidence at 15 months and these tumors exhibited a high degree of LOH^20^. *tp53* null zebrafish developed a wider range of tumors with zMPNSTs, angiosarcomas, germ cell tumors, and natural killer cell-like leukemias all throughout the body with a 37% incidence at 12 mpf^13^; while *tp53* null zebrafish that co-expressed KRAS G12D and either human *TP53* C176F or P153Δ developed embryonal rhabdomyosarcoma in the head, trunk, and tail with the C176F being a hypomorphic allele with a lower tumor incidence and the P153Δ mutation demonstrating GOF with a predisposition to head tumors^41^. However, these mutations do not represent *TP53* hotspot GOF missense mutations, which comprise the majority of *TP53* mutations in LFS patients.

As the first *tp53* “hotspot” zebrafish, R217H and R242H mutants are more analogous to the phenotypes and tumor characteristics observed in human LFS patients than previous zebrafish *tp53* mutants. A wide range of tumors developed throughout the body with histology that resembles both spindle cell-like sarcomas and SRBCs with varying histological patterns reminiscent of other sarcomas. Several of the spindle cell-like tumors were zMPNSTs, which is the most common tumor that arises in previously reported *tp53* zebrafish mutants^13,20,29,42^, and we also saw tumors similar to angiosarcomas, melanomas, leukemias, and aRMS based on histological features and previous characterizations. Of note, the *tp53* point mutants are in the *casper* background which lacks the *mitfa* gene necessary for the development of neural crest melanocytes^12^. As melanoma arises from melanocytes, the observation of melanoma in these mutants is surprising and warrants further investigation. Interestingly, we observed a different anatomical tumor distribution for each genotype with flank tumors most prevalent in *tp53* null mutants and abdominal tumors most prevalent in R217H mutants, suggesting mutation-specific mechanisms influencing tumor development. Further, we observed a sex bias in R242H mutants with predominantly females developing tumors. In LFS, the lifetime risk of developing cancer is higher in females than males, which is generally attributed to the high risk of breast cancer but this may not fully explain this finding as suggested by our results^43^.

R242H mutants have the highest lifetime tumor incidence reported out of all *tp53* mutants reported to date with a 78% incidence at 16 mpf. Interestingly, heterozygous *tp53* R217H mutants had a 42% incidence of tumor formation at 16 mpf, which is nearly the same as for the homozygotes, while heterozygous *tp53* R242H mutants had a much lower rate of 22% at 16 mpf. This suggests that the dominant negative activity of the R217H allele is sufficient for tumorigenesis, as loss of the wildtype allele has only a minor increase on tumor development whereas loss of the wildtype allele is key for aggressive tumorigenesis associated with the R242H mutation. The only other heterozygous zebrafish *tp53* mutation carriers previously reported to develop tumors is the *tp53* I166T/+ line, whereas *tp53* N168K/+, M214K/+, and *tp53* null/+ zebrafish very seldom develop tumors^13,20,29^. Of these four different mutants, only *tp53* I166T mutants exhibit dominant negative activity which likely contributed to *tp53* I166T/+ tumor development^20^.

In mice, the *Tp53* R245W (zebrafish R217W and human R248W) mutation is associated with very aggressive tumorigenesis, an increased tumor incidence^44^, and an earlier tumor onset^45^. However, the human *TP53* R248H (zebrafish R217H) mutation has not been studied nor has it been implicated in any human tumor databases. Studying this R217H mutation allowed us to decipher the impact of different amino acid changes at this residue. While *tp53* R217H mutants developed spontaneous tumors, they had a later time to tumor onset and much lower lifetime tumor incidence compared to *tp53* null and R242H mutants. This combined with the other hypomorphic characteristics of the R217H allele suggests that while the R248Q/W are considered hotspot mutations, the most common hotspot amino acid change of arginine to histidine here does not confer the aggressive tumorigenic characteristics typically associated with a hotspot mutation.

Altogether, novel GOFs in R217H and R242H mutants may account for the several month earlier tumor onset and differential tumor spectrum observed compared to other zebrafish models described in the literature. We showed that both global gene expression and DNA methylation are vastly different in *tp53* null mutants compared to R217H and R242H mutants. This suggests that there are likely major underlying mechanistic differences between complete loss of the p53 protein compared to missense p53 proteins, which needs to be considered in specific disease modeling. Of note, *tp53* null mutants are in a CG1 strain background and the R217H and R242H mutants are in a *casper* (AB strain) background, so strain variation may contribute to some observed differences.

### Transcriptomic and DNA methylome changes highlight metabolic reprogramming in mutant larvae prior to tumor formation

Having established that there are key differences in the tumor phenotypes that arise in *tp53* R217H and R242H mutants, we undertook RNA sequencing and WGBS to investigate these mutation-specific differences. GO BP and KEGG pathway analyses on DEGs and DMRs revealed broadly overlapping changes in cellular processes including metabolism, cell signaling, and the synthesis, degradation, and transport of biological macromolecules. Notably, many of the top KEGG pathways involved metabolism and biosynthesis/degradation of amino acids, including involvement of glycine, serine, and threonine metabolism in *tp53* null and R242H mutants. The reprogramming of amino acid metabolism is a well-known phenomenon in cancer as they can serve as energy sources, building blocks, and are vital for many cellular processes^46–48^. Specifically, glycine and serine are essential metabolites for cancer cells and mutant p53 has been shown to stimulate de novo serine and glycine synthesis^49,50^. Taken together, metabolic reprogramming vital to cancer cell growth and proliferation occurs in larval stages prior to tumor formation in *tp53* mutant zebrafish. Thus, characterization of the metabolome in *tp53* mutant zebrafish, both in larval stages and in tumors, will likely further elucidate tumor-driving mechanisms and potential novel therapeutic targets.

In summary, we have shown that *tp53* R217H and R242H are the most representative LFS zebrafish mutants generated to date as they recapitulate critical LFS phenotypes and affect two of the most common p53 residues found in both LFS and sporadic cancers. Going forward, these mutants can be used to shed light on the many roles of mutant p53, uncover specific cancer-driving mechanisms, and be applied to better understand complex p53 biology in many different diseases. Importantly, they present a unique preclinical platform to discover and evaluate chemopreventive therapies that can restore normal p53 function to potentially delay/prevent tumor onset for individuals with LFS.

## Supporting information

Key Resource Table

## Acknowledgments

This research was funded by the Canadian Institutes of Health Research (CIHR) (Grant #PJT-166081). KK was supported by the Ontario Graduate Scholarship (2022) and CIHR Canada Graduate Scholarship – Doctoral (2023). Flow cytometry was performed at the University of Ottawa Flow Cytometry and Virometry Core Facility (RRID:SCR_023306). Irradiation was performed at the University of Ottawa Preclinical Imaging Core Facility (RRID:SCR_021832). All DNA sanger sequencing was performed by the Ottawa Hospital Research Institute StemCore Laboratories Core Facility (RRID:SCR_012601). We would like to acknowledge Canada’s Michael Smith Genome Sciences Centre, Vancouver, Canada for the whole genome bisulfite sequencing and genome alignment. We thank Dr. David Langenau for the CG1 *tp53* null zebrafish line. We would like to thank the University of Ottawa Animal Care and Veterinary Staff for zebrafish husbandry.

## Author Contributions

**K.K:** Conceptualization, data curation, formal analysis, funding acquisition, performed experiments, methodology, validation, visualization, writing – original draft, writing – review and editing. **L.T:** Performed experiments. **E.B:** Performed experiments. **J.A.F:** Performed experiments. **K.J.C.G:** Performed experiments. **C.M:** Performed pathology review and interpretation. **B.L:** Data curation, formal analysis. **V.S:** Data curation, formal analysis. **T.T.W:** Data curation, formal analysis. **M.H:** Data curation, formal analysis, supervision. **M.M:** Data curation, formal analysis, investigation. **A.C:** Data curation, formal analysis, investigation. **Q.C:** Performed experiments. **A.S:** Data curation, formal analysis, supervision. **D.M:** Conceptualization, data curation, formal analysis, supervision. **S.V.P:** Conceptualization, data curation, formal analysis, funding acquisition, performed experiments, methodology, validation, visualization, supervision, writing – original draft, writing – review and editing. **J.N.B:** Conceptualization, funding acquisition, supervision, writing – original draft, writing – review and editing.

## Materials and Methods

### Zebrafish Husbandry

*Danio rerio* (zebrafish) lines (*tp53* null – CG1 background, *casper, tp53* R217H, and R242H – AB background) were maintained at 28°C on a recirculating water system with a 14:10 hour light:dark cycle at the University of Ottawa Aquatics Facility. Larval animals were fed rotifers and Gemma75 and juvenile animals were fed Gemma150 four times a day while adult animals were fed Gemma300 twice a day (Gemma Micro, Skretting). Embryos were produced from both single and group crosses and were maintained at 28°C^51^ in E3 medium (5 mM NaCl, 0.17 mM KCl, 0.33 mM CaCl_2_, 0.33 mM MgSO_4_) and were maintained in approx. equal sex distributions until 18 mpf. Embryos were staged according to standard practice^52^. Animal use in this study was approved by and conducted in accordance with the University of Ottawa’s Animal Care Committee under protocols #4166 and #4239 which are in line with the Canadian Council for Animal Care guidelines.

### Generation of CRISPR Zebrafish Mutants

Generation of the *tp53* R217H point mutant was previously described in detail^11^ and the R242H allele was done similarly as follows. Cas9 mRNA was made from pT3TS-nCas9n plasmid after its linearization with XbaI using mMessage mMachine T3 kit (Thermo Fisher Scientific, AM1348) and purified with LiCl precipitation according to the kit instructions. The pT3TS-nCas9n was a gift from Wenbiao Chen (Addgene plasmid #46757; RRID:Addgene_46757)^53^. The sgRNA template for *tp53* R242H knock-in was generated by performing an overlap-extension PCR of p53-R242_sgRNA-2_sense and Rev-sgRNA-scaffold oligos using Taq DNA polymerase (ABM, G009) by combining 10 μl of 10× buffer, 6 μl of 25 mM MgSO4, 2 μl 10 mM dNTP, 5 μl of each oligo at 25 μM, 71 μl water and 1.5 μl of Taq and running the following program: 94°C for 5 min; 5 cycles: 94°C for 30 s, 55°C for 30 s, 72°C for 30 s. The resulting PCR products were purified and in vitro transcribed using MEGAshortscript T7 kit (Thermo Fisher Scientific, AM1354). The sgRNA was purified according to the kit instructions. Injections were performed with Cas9 mRNA at 300 ng/μl, sgRNA at 150 ng/μl and the single-stranded oligo R242H-S-Asymm_oligo at 1 μM into 1-cell zebrafish embryos. Assessment of sgRNA efficiencies was performed using either T7 Endonuclease I (NEB, M0302S) digestion according to the manufacturer’s protocol or using heteroduplex mobility assay^54^.

DNA for genotyping was prepared as previously described^55^. PCRs were run with appropriate primers (Supplementary Table S2) using *Taq* DNA polymerase following the touch-down PCR protocol: 94°C for 3 min; 10 cycles: 94°C for 30 s, T_a_°C + 10 (with 1°C decrease every cycle), 72°C for 30 s, 25 cycles: 94°C for 30 s, T_a_°C, 72°C for 30 s. R242H knock-in injected embryos and potential F0 and F1 adults were genotyped using allele-specific PCR assays with either *R242H-KI_for* (R242H allele) or *R242-WT_for* (wild-type) and R242H_SA_R primers using the above program with T_a_=51. Primers to genotype stable mutants are R217H_SA (T_a_=55) or R242H_SA (T_a_=53). PCR products were enzymatically digested (Supplementary Table S2) for 2 hours prior to gel electrophoresis: 7μL PCR product, 0.5μL enzyme, 0.2μL 10X Cutsmart buffer, and 10.5μL nuclease free water. The final PCR products were visualized using standard 2% agarose gel electrophoresis.

### Embryo Dechorionation

To remove chorions, embryos <3 dpf were bathed in 0.05mg/mL Pronase (Sigma, 11459643001) for 15-20 minutes at RT until all embryos came out of their chorions (either naturally or with gentle pipetting) followed by 3X washes in E3 medium.

### X-Ray Irradiation

An X-RAD 320 (320kV, 10mA; Precision X-Ray Inc) was used to irradiate embryos to induce DNA damage and activate the p53 pathway. Embryos were contained in 15mL of E3 in 10cm Petri dishes placed 50cm from the X-Ray source. X-Rays passed through the F1 filter (2mm Al, Precision X-Ray Inc) and embryos were dosed with either 16 Gy (123328 MU) or 30 Gy (231240 MU).

### Camptothecin Treatment

Camptothecin (CPT) (Sigma, C9911) (0.1μM) was prepared fresh from 2mM stock in DMSO (stored at −20°C) to methylene blue-free E3 medium. 24 hpf dechorionated embryos were placed in 12-well plates and bathed in 0.1μM CPT or DMSO for 4 hours at 28°C. Following treatment, embryos were washed 3X with methylene blue-free E3.

### Acridine Orange Staining

To assess apoptosis, 28-30 hpf dechorionated embryos were bathed in 2 µg/ml AO (ThermoFisher, A1301) for 30 mins and subsequently washed 3X with methylene blue-free E3. Fish were anesthetized (MS-222) and imaged immediately using the FITC channel (Filter set 90 HE LED (E), LED 475) on a ZEISS inverted Axio Observer 7 microscope equipped with a Colibri 5/7 illumination system and a ZEISS Axiocam 705 mono CMOS camera (Carl Zeiss Microimaging Inc.). Image visualization and processing was done in Fiji (RRID:SCR_002285)^56^ and ZEISS ZEN Microscopy Software (RRID:SCR_013672).

### Morpholino Knockdown

An mdm2 splice-blocking MO (5′-TGTTAAGAGATTCAGTACGCACCGC-3′)^20^ (ZDB-MRPHLNO-100429-9) at a concentration of 0.7mM was injected into the yolk of one-cell stage embryos as a bolus of approximately 20% of the yolk. A final concentration 0.5% Phenol red was added to visualize the injection solution. Injection parameters were in accordance with standard protocols^57,58^. Larvae were imaged at 72 hpf on a ZEISS Axio Zoom.V16 microscope and a ZEISS Axiocam 208 color camera (Carl Zeiss Microimaging Inc.). To verify if splicing occurred, PCRs were run with the *mdm2_MO1* (T_a_=51) and *mdm2_MO2* (T_a_=51) primers (Supplementary Table S2) using *Taq* DNA polymerase following the touch-down PCR protocol: 94°C for 3 min; 10 cycles: 94°C for 30 s, T_a_°C + 10 (with 1°C decrease every cycle), 72°C for 30 s, 25 cycles: 94°C for 30 s, T_a_°C, 72°C for 30 s. The final PCR products were visualized using standard 2% agarose gel electrophoresis.

### Embryo Protein Extraction and Western Blotting

Approximately fifty 30 hpf dechorionated embryos (+/- 30 Gy 6 hpi) were collected into a 1.5mL Eppendorf tube and placed on ice for several minutes. Embryos were rinsed three times with cold Ringer’s solution (116mM NaCl, 2.9mM KCl, 5.0mM HEPES, pH 7.2), deyolked by pipetting up and down twelve times in deyolking buffer (0.3mM PMSF (phenylmethylsulfonylfluoride) and 1mM EDTA in Ringer’s solution) and rinsed twice with cold Ringer’s solution. Samples were placed in 300μL of the lysis buffer (1X Ripa Buffer (Sigma, 20-188), 1X cOmplete™ Mini Protease Inhibitor Cocktail (Roche, 04693159001), 2mM sodium orthovanadate, 10mM sodium fluoride, 20mM sodium beta-glyercolphosphate, and 1mM PMSF) and lysed by passing solution through a 23G needle and 1mL syringe twelve times. Tubes were centrifuged at 4°C at full speed for 3 minutes and supernatant was transferred to a new tube, with 10μL saved for the BCA assay. 100μL of 4X Laemlli loading buffer (Bio-Rad, #1610747) with β-mercaptoethanol was added and each sample was boiled at 95°C for 5 minutes. The protein concentration was quantified using the Pierce™ BCA Protein Assay Kit (Thermofisher, #23227). Samples were run on a stain free acrylamide gel (Bio-Rad, #1610183) at 200V for 30 minutes and gels were activated on a UVP ChemStudio touch Imaging System (Analytik Jena, SP-1082) for 1-5 minutes. The gel was transferred to an Immun-Blot® Low Fluorescence PVDF membrane (Bio-Rad, 1620261) at 100V for 45 minutes. The blot was blocked in a 5% skim milk/1X TBS-T (Tris-Buffered Saline, 0.1% Tween-20) (ThermoFisher, J60764.K3) (Sigma, P2287) solution for 1 hr shaking at room temperature (RT). The p53 protein was detected by hybridization with the mouse ZFp53-9.1 primary antibody (Abcam Cat# ab77813, RRID:AB_10864112) at 1:500 in 2.5% skim milk solution overnight shaking at 4°C which detects a band of approximately 53 kDa^59^. The blot was washed with TBS-T 4X 10 mins shaking at RT and then was incubated with the Goat anti-Mouse IgG (H+L) Cross-Adsorbed Secondary Antibody, HRP (ThermoFisher, G-21040, RRID:AB_2536527) at 1:1000 in 5% skim milk solution for 1 hr shaking at RT. The blot was washed with 1X TBS-T 3X for 10 mins shaking at RT and then washed with 1X TBS for 10 mins at RT. The blot was imaged on a UVP ChemStudio touch Imaging System (Analytik Jena, SP-1082) prior to signal detection to capture total protein amount per lane. Signals were detected using the SuperSignal™ West Dura Ultimate Sensitivity Substrate (ThermoFisher, 37071) and imaged on the UVP ChemStudio. Band weights were determined using the Precision Plus Protein™ All Blue Prestained Protein Standards (Bio-Rad, #1610373).

### RNA Extraction and cDNA Synthesis

Each RNA sample was extracted from fifty zebrafish embryos anesthetized with tricaine (MS-222) and homogenized in 500μL TRIzol reagent (Life Technologies, 15596026) using a 1mL syringe and 23G needle. Lysed samples were purified and DNAse treated using the Direct-zol RNA MicroPrep kit (Zymo Research Corporation, R2060). cDNA was synthesized by adding 2μg of total DNAse-treated RNA to 2μl of 100μM oligo-dT(15-18) (Integrated DNA Technologies) and heating at 70°C for 10 min before cooling on ice. Next 2μl of M-MuLV buffer (NEB, M0253S), 0.25μl of Protector RNase Inhibitor (Roche, 3335399001), 0.25μL of M-MuLV Reverse Transcriptase (New England BioLabs, M0253L) and 1.6μL of water were added to the reaction and incubated at 42°C for 1 hour followed by heating at 90°C for 10 min before cooling.

### Quantitative PCR

Quantitative PCR was performed on a Bio-Rad C1000 Thermal Touch CFX96 and CFX384 Real-Time Systems (Bio-Rad) using BlasTaq™ 2X qPCR MasterMix (ABM, G891) and cDNA was diluted to 1:40. qPCR cycling conditions were as follows: 95°C for 3 minutes, [95°C for 10 s, 55°C for 10 s, 72°C for 30 s] (40 cycles) with primers listed in Supplementary Table S2. *β-actin* and *elfa*^60^ were used as housekeeping genes and expression values were calculated using the CFX Maestro Software (v2.3) (Bio-Rad). Fold changes were calculated by normalizing data to either the DMSO or 0 Gy *casper* control samples for each gene, respective to treatment group. A Grubbs’ test was performed to identify and exclude outliers using an alpha <0.5. Statistical significance was calculated with a two-way ANOVA and Tukey’s Honest Significant Difference test.

### Whole Mount Immunofluorescence

Dechorionated embryos were euthanized (MS-222, 1mg/mL) and fixed overnight at 4°C in 4% paraformaldehyde (PFA) (Sigma, P6148) in 1X PBS (ThermoFisher, AM9625). Embryos were washed twice in 1X PBS for 5 minutes, once in deionized water for 5 minutes, and blocked for 1 hour in blocking solution (1X PBS, 0.01% bovine serine albumin (Fisher, SH3057402), 0.02% sheep serum (Sigma, S22), 0.1% Triton X-100, 0.01% DMSO). Embryos were incubated in the primary antibody, rabbit IgG, anti-phospho-Histone H3 (Ser10) (diluted 1:500 in blocking solution, Millipore Sigma, 06-570, RRID:AB_310177) overnight rocking at 4°C. Subsequently, embryos were washed for 8 hours in 0.1% Triton X-100/1X PBS (with solution changes several times throughout the day) and incubated overnight at 4°C in the dark in the Goat anti-Rabbit IgG (H+L) Cross-Adsorbed Secondary Antibody, Alexa Fluor™ 488 (diluted 1:750 in blocking solution, Thermofisher, A-11008, RRID:AB_143165). Embryos were rinsed 2 x 5 minutes in 0.1% Triton X-100/1X PBS and were stored in 1X PBS at 4°C until imaged (maximum two weeks before imaging). Stained zebrafish embryos were imaged using the FITC channel (Filter set 90 HE LED (E), LED 475) on a ZEISS inverted Axio Observer 7 microscope equipped with a Colibri 5/7 illumination system and a ZEISS Axiocam 705 mono CMOS camera (Carl Zeiss Microimaging Inc.).

### Flow Cytometry

Samples for flow cytometry were collected by placing 20-25 dechorionated embryos in an Eppendorf tube on ice. Embryos were deyolked in 180μL cold Ringer’s solution (116mM NaCl, 2.9mM KCl, 5.0mM HEPES, pH 7.2) by pipetting up and down three times. Samples were rinsed twice with 1X PBS and dissociated at 35°C for 5-10 minutes with a 5.4 mg/mL collagenase/PBS solution (Sigma, C0130). Next, 20% fetal bovine serum (Wisent, 080-450) was added to supernatant to terminate the enzymatic reaction once dissociation was complete. The samples were centrifuged (300xG at 4°C for 5 minutes), the supernatant was removed, and the cells were resuspended in 500μL 4% PFA/PBS for 15 minutes at RT and then 500μL of 1X PBS was added to the tubes. The samples were centrifuged (600xG at RT for 5 minutes) and were rinsed in 1X PBS, centrifuged, and the pellet was resuspended in a 0.1% Triton X-100/0.375% sodium citrate/PBS solution for 15 minutes at RT. Samples were centrifuged again, resuspended in 400μL 1X PBS and filtered through a 35μm strainer. To stain DNA, 1 drop of FxCycle™ Violet Ready Flow™ Reagent (Thermofisher, R37166) was added and samples were run on an Aurora Flow Cytometer (Cytek Biosciences). The flow cytometry results were analyzed using Kaluza Analysis Software (Beckman Coulter). Cells were gated to remove debris and doublets and limited to the first 25000 (+/-100) cells for each sample (Supplementary Figure S8).

### Tumor Watch Analysis

The tumor watch experiment followed tumor development in *tp53* null, R217H, R242H, adult zebrafish beginning at 6 mpf and ended at 16 mpf. The animals were frequently monitored for any signs of tumor development, and tumor-bearing fish were euthanized for histological analysis. Additionally, animals were counted every two weeks and tumor-free survival was calculated based on number of surviving fish compared to starting number for each time interval. Survival analysis was performed using R with ‘survival’, ‘survminer’, ‘Rcpp’, and ‘ggplot2’ and the survdiff function was used to determine the statistical significance^61^.

### Histology

Adult zebrafish were fixed in 10% neutral buffered formalin (NBF) (Sigma, HT501128) for a minimum of ten days, bisected in the sagittal plane and processed whole, using a Leica ASP6025S tissue processor. The processing steps include: 2 hrs in NBF, 1 hr in NBF, 45 mins in 70% ethanol (EtOH), 45 mins in 95% EtOH X2, 45 mins in 100% EtOH X3, 45 mins in xylene X2, 30 mins in Paraplast Surgipath paraffin (Leica, 39601006) X2, and 40 mins in paraffin X2. Next, fish were embedded in paraffin and sectioned on a Leica RM 2235 microtome. Slides were stained on a Somagen Tissue-Tek Prisma using standard hematoxylin and eosin and coverslipped by a Somagen Tissue-Tek Glas. Slides were examined by a board-certified pediatric and anatomical pathologist (C.M.) using a Nikon Eclipse Ni light microscope. Digital images were captured with a 5-megapixel Olympus SC50 microscope mounted camera using Olympus cellSens software. zMPNST tumors were characterized based on three main features: a spindle cell component (can be variable, contains varying amounts of small round cells, and usually termed “epithelioid” in human tumor literature); a storiform/whorled/fascicular growth pattern; and the presence of clefts.

### Loss of Heterozygosity Genotyping Strategy

The genotyping strategy involved amplifying a ∼300bp genomic region around the point mutation with PCR and then performing a restriction enzyme digest where the restriction enzyme recognizes the point mutation and cuts the product in half, producing one smaller band which is then visualized using agarose electrophoresis (as described in the point mutant generation section). Wildtypes have one larger band, heterozygous mutants have two bands (large and small), and homozygous mutants have a predominantly smaller band (Supplementary Figure S5A). Complete digestion with restriction enzymes does not always occur, so often the heterozygous mutants have a larger upper band while homozygous mutants have a significant lower band.

### RNA Sequencing and Analysis

Total RNA samples were processed as standard RNA-seq by Azenta Life Sciences. The RNA-Seq data were mapped with STAR (v2.7) (RRID:SCR_004463)^62^ to generate BAM files and transcripts were quantified with RSEM (RRID:SCR_000262)^63^ using the zebrafish V4.3.2 transcript annotation^64^. Differentially expressed genes were identified using the ‘DESeq2’ package (1.42.0) (RRID:SCR_015687) in R and genes were filtered to remove those with counts of less than 10 in at least three samples^65^. For hypothesis testing we applied the Wald test and ashr shrinkage^66^. Differentially expressed genes were identified as those with a fold change of ±2 and a p-value of less than 0.05 as parameters. Variance stabilizing transformation was performed as a function of the DESeq2 library^65^. Heatmaps were produced using the ‘ComplexHeatmap’ package and upset plots were created using the ‘ComplexUpset’ package. Known p53 targets from IARC (International Agency for Research on Cancer) were mapped to zebrafish orthologues and these were intersected with the differentially expressed genes to create the heatmap of p53 target genes. FishEnrichr^67–69^ was used for GO enrichment of biology processes (2018) and KEGG pathway analysis (2019) (RRID:SCR_012773) by analyzing the differently expressed genes and their corresponding adjusted p-value for each comparison. The top ten terms were ranked using the combined score and were visualized with dotplots produced with the ‘stringr’ and ‘forcats’ packages in R.

### WGBS Sequencing and Analysis

Thirty pooled 5 dpf larvae were flash frozen in an Eppendorf tube, then gDNA extraction and methylation libraries were generated at the Michael Smith Laboratories, University of British Colombia, Vancouver, Canada using the Post-Bisulfite Adapter Ligation method as published^70^, except the starting material is 100ng of gDNA from bulk samples, only one round DNA generation/random priming was used, PCR was limited to four rounds, and after PCR a size selection step enriched for larger fragments of DNA. Uniquely indexed libraries were pooled and submitted for sequencing to Michael Smith Genome Sequencing Centre, Vancouver, Canada on the NovaSeq6000. Alignment and fractional methylation calls were generated using GemBS at Michael Smith Laboratories, University of British Columba, Vancouver, Canada.

We used methylKit (RRID:SCR_005177) to evaluate the zebrafish methylation data^71^. To preprocess the data, we filtered out regions with less than 10 reads and the highest 0.1% coverage regions as recommended by methylKit. We also normalized coverage between samples. We binned across the genome in 1,000bp bins with a step size of 1,000bp and used Fisher’s Exact test to determine differentially methylated bins between the groups. To determine which genotypes were significantly different for each region, we conducted pairwise logistic regression tests for the differentially methylated regions previously found. For both the Fisher’s Exact tests and logistic regressions, we corrected for multiple hypothesis testing with the Benjamini Hochberg method and used a false discovery rate cut-off of 0.05 and a methylation difference cut-off of 25%.

We annotated the significantly methylated regions and applied FishEnrichr for GO enrichment of BP (2018) and KEGG pathway analysis (2019) and terms were ranked using the combined score^67–69^. The top ten terms for each comparison were visualized with dotplots produced with the ‘stringr’ and ‘forcats’ packages in R.

### Statistics and Data Visualization

Data were collected within Microsoft Excel (Office 365 for Mac, Microsoft) (RRID:SCR_016137) and imported into R Studio (Version 4.0.0) for visualization and statistical analysis. The ‘ggplot2’, ‘magrittr’, ‘ggpubr’, ‘ggsci’, ‘tidyverse’, ‘matrixStats’, ‘broom’, ‘AICcmodavg’, ‘dplyr’, ‘tidyr’, ‘scales’, ‘Cairo’, and ‘readxl’ packages were used within R software to process data and generate plots. Unless otherwise specified, error bars represent standard deviation and experiments were performed three times. p-values are indicated in the figure legends.

## Supplemental Information

Document S1. Figures S1-S8 and Tables S1, S2

