## Supplementary material for "*tp53* R217H and R242H Mutant Zebrafish Exhibit Dysfunctional p53 Hallmarks and Recapitulate Li-Fraumeni Syndrome Phenotypes": Document S1

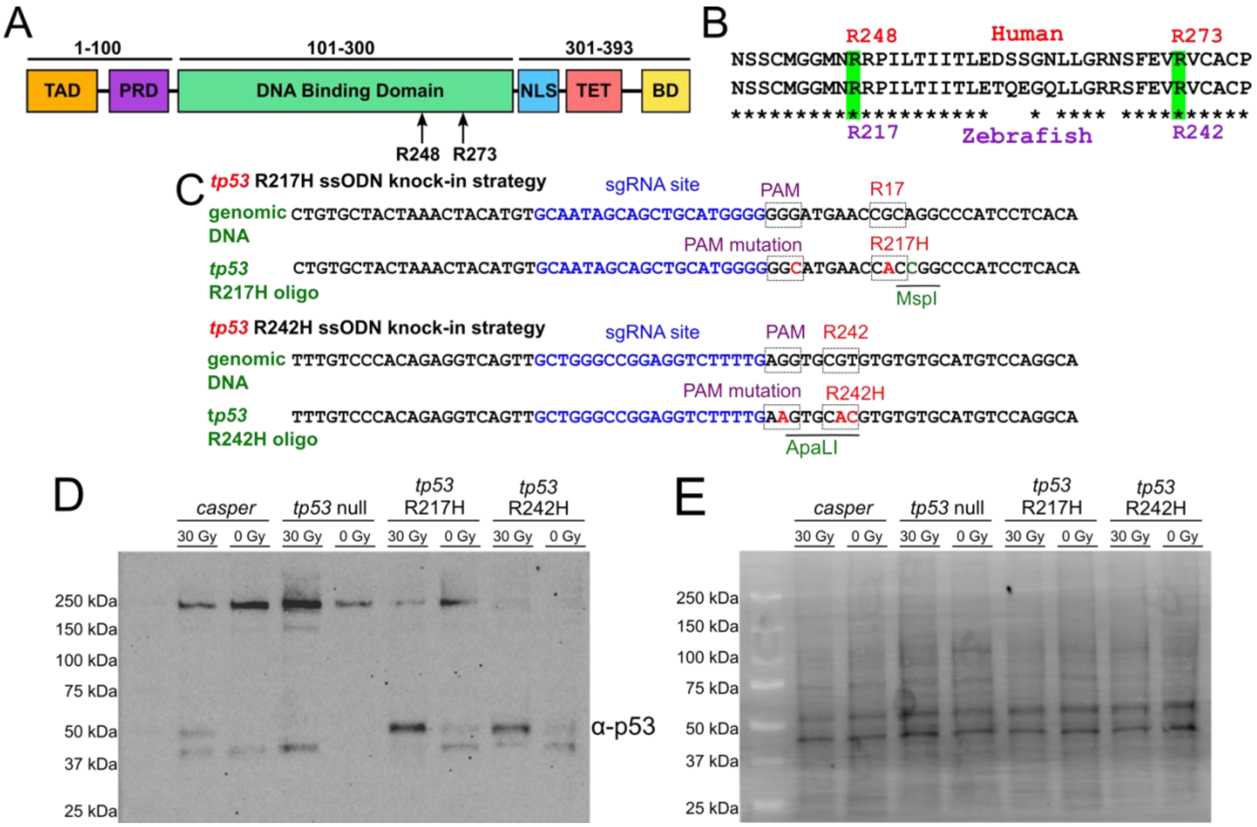


**Supplemental Figure 1. Generation of *tp53* R217H and R242H CRISPR knock-in mutants which both display increased p53 protein levels following IR.** (A) Schematic showing the human p53 protein domain structures including the transactivation domain (TAD), proline rich domain (PRD), DNA binding domain, nuclear location signal area (NLS), tetramerization domain (TET), and the basic domain (BD). Arrows indicate the location of the R248 and R273 hotspot mutations in the DNA binding domain. (B) Alignment of the human and zebrafish p53 amino acids showing the high conservation in the area surrounding the human R248 and R273 (zebrafish R217 and R242) residues. Conserved amino acids are indicated by a "*". (C) CRISPR ssODN (single-stranded oligodeoxynucleotides) knock-in strategies to create the zebrafish p53 R217H and R242H mutants. (D) Representative western blot for α-p53 (~53 kDa) from 50 pooled 30 hours post-fertilization (hpf) embryos treated with +/- 30Gy of IR collected 6 hours post-IR (hpi), n=4 replicates. (E) Stain-free blot showing the total protein per sample as a loading control.

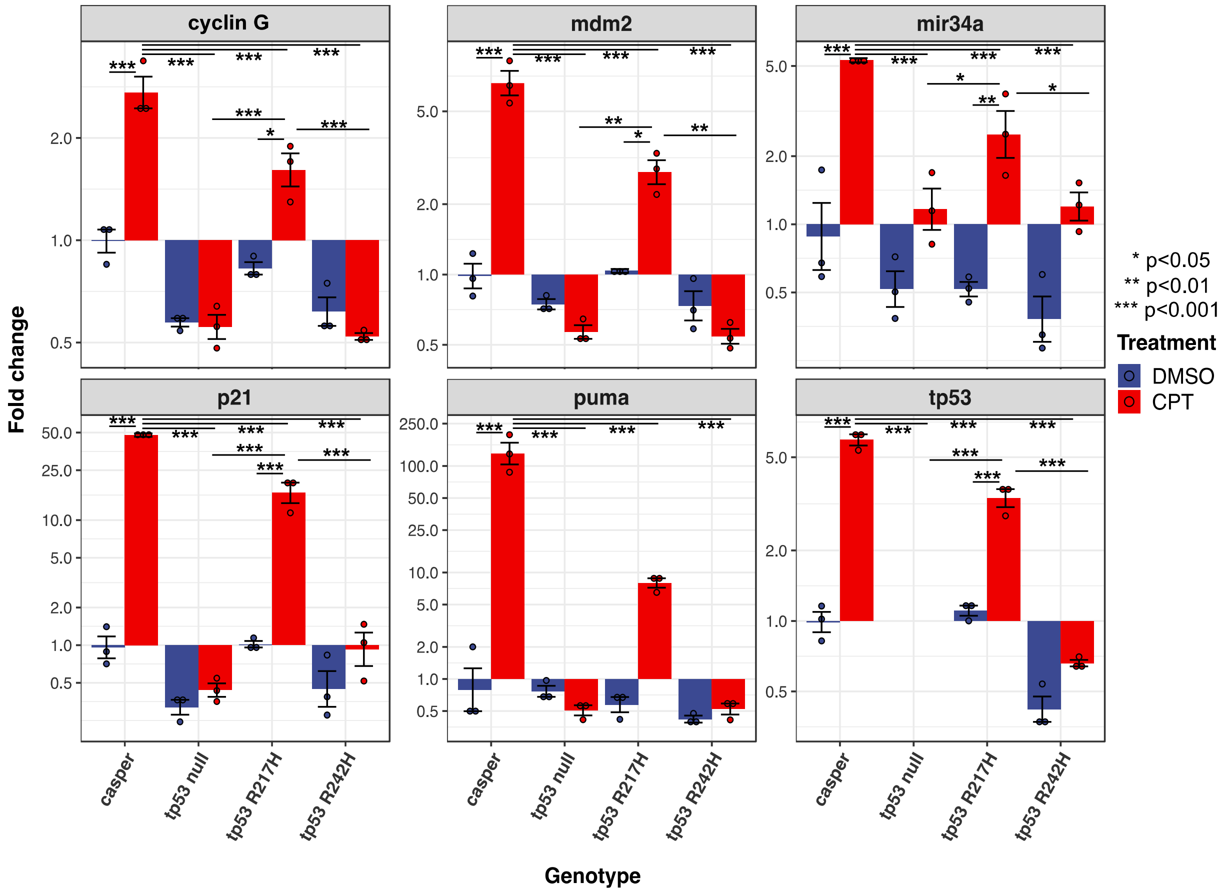


**Supplemental Figure 2. CPT treatment decreases the fold change of expression of several p53 target genes in *tp53* mutants.** Twenty-five embryos per sample were collected at 30 hpf following a 4-hour treatment with 0.1µm Camptothecin (CPT) or DMSO. Expression of *tp53* was not detected in the *tp53* null samples. Data represents two technical replicates and three biological replicates. The fold change of each gene was normalized to the *casper* DMSO treated group. Statistical significance was determined by a two-way ANOVA and a Tukey's Honest Significant Difference test. Error bars represent standard deviation. The Y-axis is logarithmic scale 10.

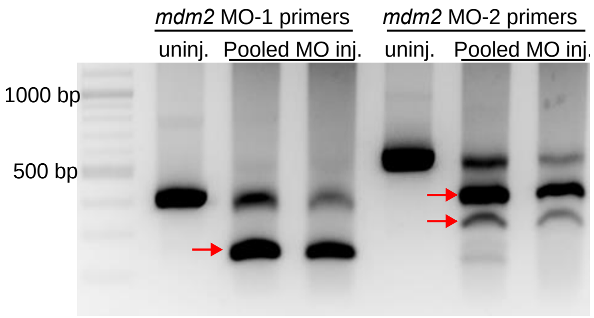


**Supplemental Figure 3. *mdm2* MO leads to incorrect splicing of *mdm2* mRNA.** Agarose gel imaging showing the *mdm2* MO PCR that verified that MO injection resulted in incorrect *mdm2* splicing as indicated by red arrows in samples from pooled MO injected (inj.) fish and was not present in pooled uninjected (uninj) fish.

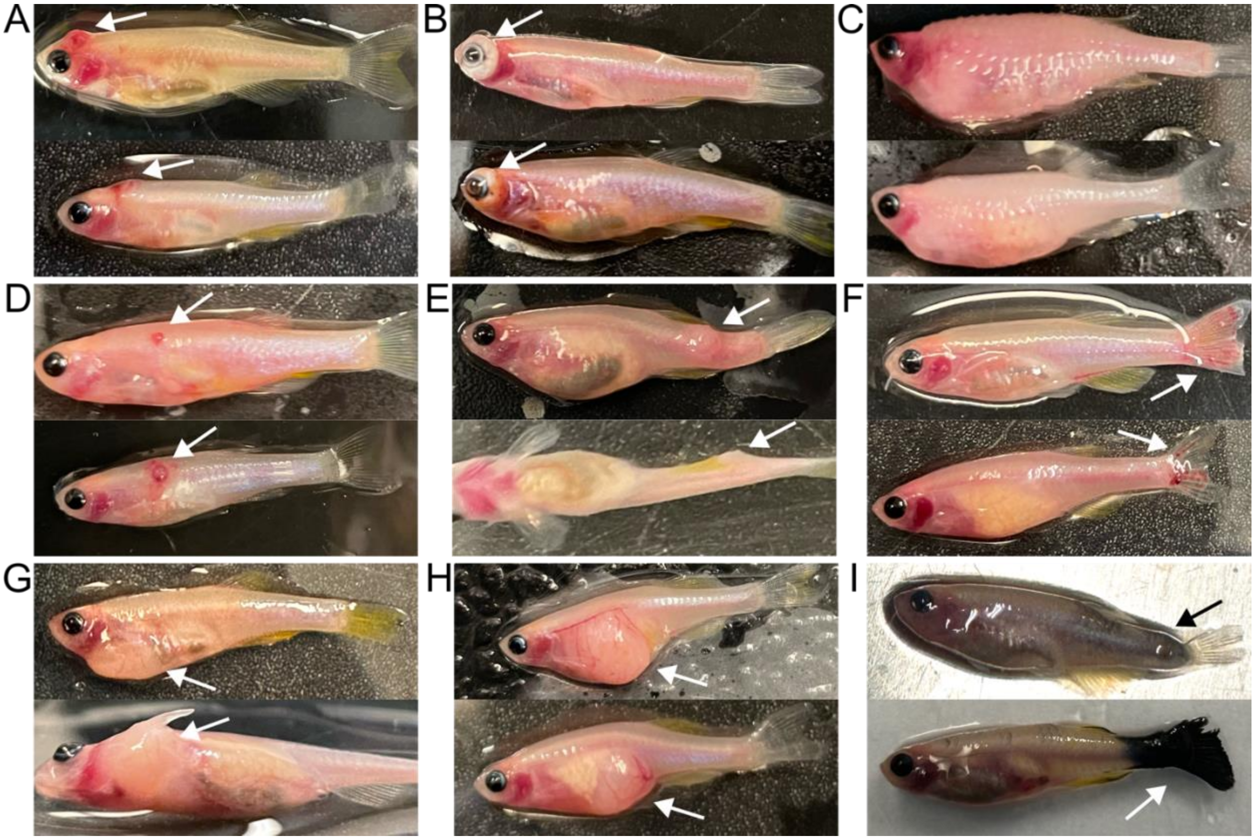


**Supplemental Figure 4. Locations of tumors that arise in *tp53* mutant fish.** Images of the different tumor locations include (A) head tumors (likely angiosarcomas); (B) eye tumors; (C) "infection" tumors as shown by the severe swelling, paleness, and protruding scales; (D) bloody flank tumors (likely angiosarcomas); (E) flank tumors (deeper in the muscle); (F) fin tumors; (G) gill tumors; (H) abdominal tumors; and (I) pigmented flank skin and tail (potentially melanoma). Arrows point to tumors.

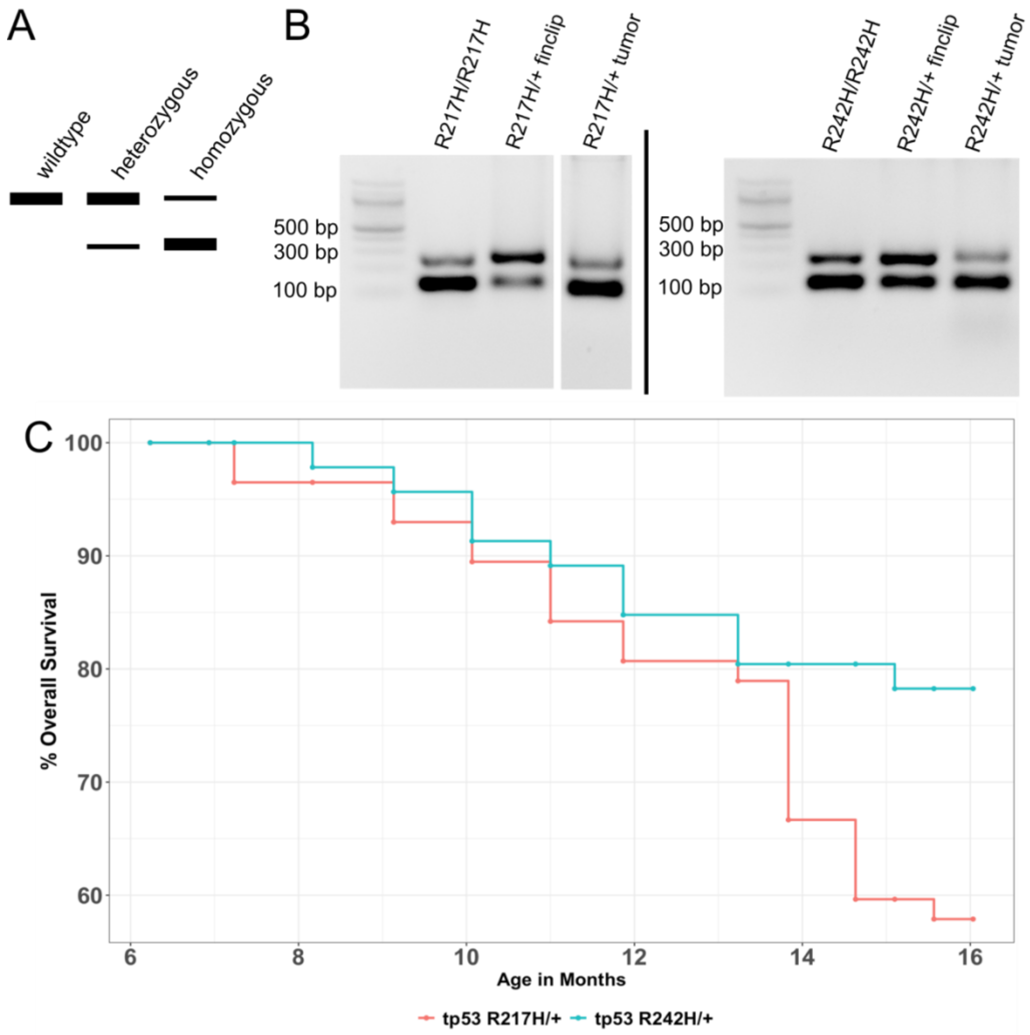


**Supplemental Figure 5. Heterozygous *tp53* point mutant tumors display loss of heterozygosity.** (A) Schematic of the *tp53* point mutant restriction enzyme genotyping assay. If the point mutation is present, the restriction enzyme will cut the PCR product, resulting in a lower band. Wildtype samples have no cutting, heterozygous mutants have a thicker upper band and faint to equal lower band, and homozygous mutants have a thicker bottom band with a minimal-to-no upper band. (B) Agarose gel images showing homozygous point mutant DNA with a paired fin-clip and tumor DNA sample from a heterozygous point mutant showing loss of heterozygosity in the tumors. The R217H/R217H DNA sample was ran on a separate gel from the R217H/+ samples. (C) Kaplan-Meier analysis showing tumor-free survival and onset in the heterozygous *tp53* R217H/+ (n=57), and *tp53* R242H/+ (n=46). Percent tumor-free survival refers to the age a specific fish had died or was euthanized due to tumor development. Tumor development was not observed in the *casper* line during this time period.

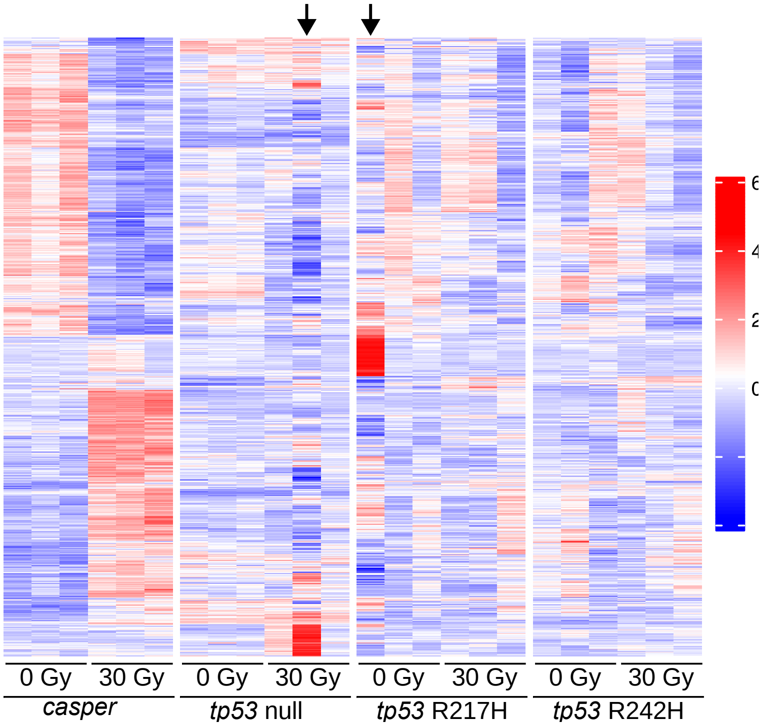


**Supplemental Figure 6. The removal of a 30 Gy *tp53* null sample and a 0 Gy *tp53* R217H sample as two technical outliers.** Heatmap showing differently expressed genes (DEGs) from all genotype comparisons from the 0 Gy treatment groups (adjusted p-value <0.05, fold change >2). Black arrows point to two samples that were removed as technical outliers due to the presence of strong upregulation of a subset of genes and vastly different expression compared to the other two biological replicates.

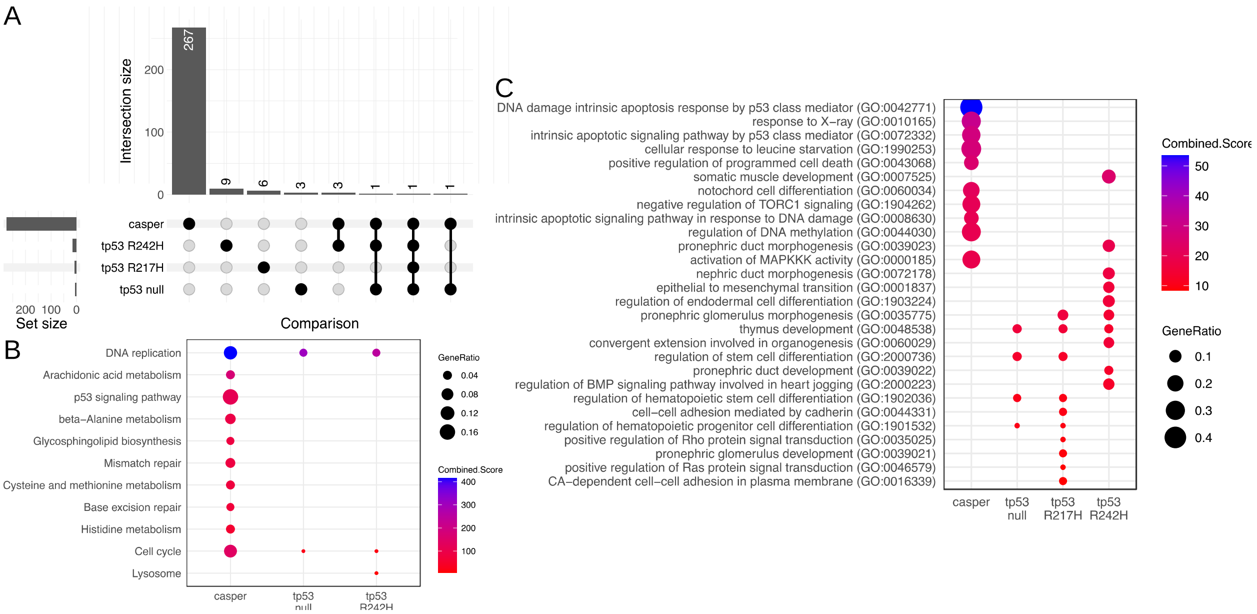


**Supplemental Figure 7. IR treatment activates the p53 pathway and apoptosis signalling pathways in *casper* but not mutant *tp53* embryos.** (A) UpSet plot shows the number and overlap of DEGs between the 0 and 30 Gy treatment groups for each genotype. (B, C) Dot plots showing the top ten KEGG pathway (B) and GO BP (C) terms, ranked by combined score, between the 0 and 30 Gy treatment groups for each genotype. Dot size corresponds to the ratio of DEGs per total number of genes for each term and the colour corresponds to the combined score. There were no KEGG pathway terms identified for the *tp53* R217H comparison.

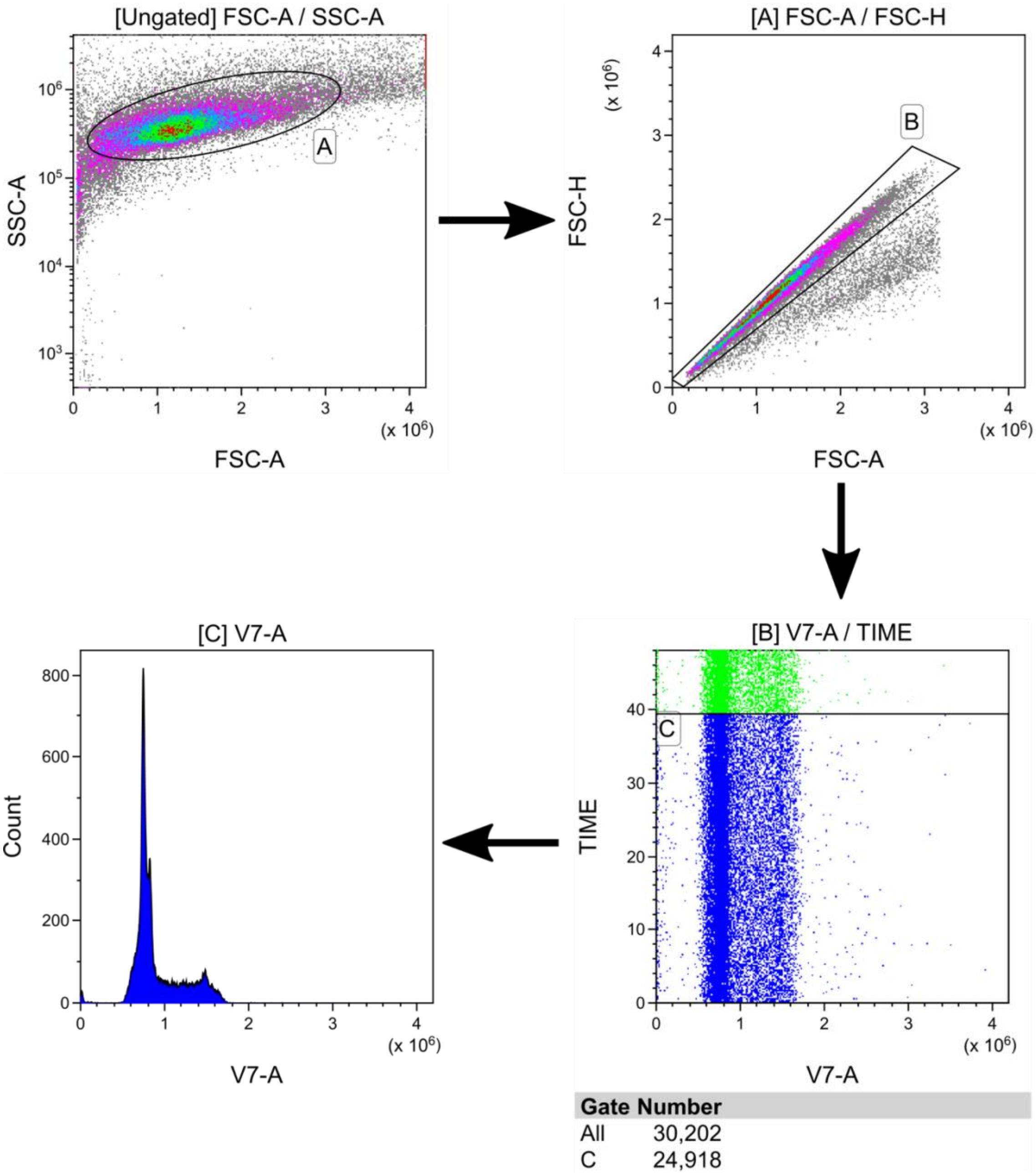


**Supplemental Figure 8. Gating strategy for flow cytometry of zebrafish embryo single cell suspensions stained with FxCycle violet.** Gate A was applied to remove debris and gate B was applied to remove any doublets. Gate C was applied to limit each sample to 25000 (+/- 100) cells to compare DNA content histograms between each sample.

**Supplemental Table 1. Age, physical location, and notes regarding histology of *tp53* mutant tumors.**

| Fish ID, genotype, and sex | Age (mpf) | Physical location | Notes |
|---|---|---|---|
| R242H 1 Male | 11.3 | Abdomen | -spindled to epithelioid cells, highly vascular with abundant capillary sized vessels, marked nuclear pleomorphism<br>-infiltrative margins with no capsule<br>-like R242H 5, R242H 6 |
| R242H 2 Female | 11.3 | Abdomen | -spindle cells, storiform pattern with clefts, c/w zMPNST pattern<br>-partial capsule or pseudocapsule<br>-partially infiltrative<br>-mild nuclear pleomorphism |
| R242H 3 Female | 11.7 | Head | -zMPNST pattern with areas of increased cellularity and SRBC morphology<br>-infiltrative margins<br>-mild nuclear pleomorphism |
| R242H 4 Male | 7 | Eye | -zMPSNT pattern with spindle cells, storiform pattern, clefts<br>-largely confined to eye<br>-mild nuclear pleomorphism |
| R242H 5 Male | 8.7 | Head | -infiltrative, eroding through skull, into eye<br>-spindled to epithelioid cells, highly vascular with abundant capillary sized vessels, marked nuclear pleomorphism<br>-like R242H 1, R242H 6 |
| R242H 6 Male | 11.3 | Flank | -spindled to epithelioid cells, highly vascular with abundant capillary sized vessels, moderate nuclear pleomorphism<br>-infiltrative margins with no capsule<br>-like R242H 1, R242H 5 |
| tp53 -/- 1 Male | 10.4 | Eye | -zMPSNT pattern with spindle cells and SRBC component, storiform pattern, clefts<br>-largely confined to eye<br>-mild to moderate nuclear pleomorphism |
| tp53-/- 2 Male | 12.4 | Abdomen | -zMPSNT pattern with spindle cells, storiform pattern, clefts<br>-infiltrative margins<br>-minimal nuclear pleomorphism |
| tp53 -/- 3 Female | 14.3 | Abdomen | -zMPSNT pattern with spindle cells and prominent SRBC component, storiform pattern, clefts<br>-infiltrative margins<br>-mild nuclear pleomorphism |
| tp53 -/- 4 Female | 13.1 | Flank | -zMPSNT pattern with spindle cells, storiform pattern, clefts<br>-infiltrative margins<br>-minimal nuclear pleomorphism |
| R217H 1 Male | 10.8 | Abdomen | -spindle cells arranged in interlacing fascicles and occasional giant cells, no clefts<br>-resembles fibroblastic or smooth muscle tumor<br>-infiltrative margins |

| | | | |
|---|---|---|---|
| | | | -mild nuclear pleomorphism<br>-unique pattern |
| R217H 2 Male | 10.8 | Eye | -zMPSNT pattern with spindle cells and SRBC component, storiform pattern, clefts<br>-largely confined to eye<br>-mild to moderate nuclear pleomorphism |
| R217H 3 Male | 13 | Abdomen | -zMPSNT pattern with spindle cells, storiform pattern, clefts<br>-infiltrative margins<br>-minimal nuclear pleomorphism |
| R217H 4 Male | 13 | Gill | -SRBC pattern with rosettes and follicle-like structures<br>-areas reminiscent of pediatric "blastomas", e.g. nephroblastoma, retinoblastoma<br>-follicle-like structures appear to contain eosinophilic material much like thyroid colloid, imparting an organoid or organ-like appearance<br>-associated with the gills/mouth, infiltrative<br>-unique tumor among this group, with others being zMPNST-like or highly vascular |
| R217H 5 Male | 10 | Flank | -diffuse, pattern-less proliferation of srbc's with interspersed giant cells<br>-another unique pattern among this group – very much resembles solid variant of alveolar rhabdomyosarcoma<br>-arising within skeletal muscle<br>-infiltrative margins<br>-mild to moderate nuclear pleomorphism |

**Supplemental Table 2. Oligos sequences and applicable restriction enzymes for digest.**

| Name | Sequence | Restriction Enzyme for Digest (If applicable) |
|---|---|---|
| p21_qfor | AGCTGCATTCGTCTCGTAGC | |
| p21_qrev | TGAGAACTTACTGGCAGCTTCA | |
| mir34a_qfor | CTGCTGTGAGTGGTTCTCTGG | |
| mir34a_qrev | GCGGCAGTATACTTGCTGATT | |
| cyclinG1_qfor | CATCTCTAAAAGAGGCTCTAGATGG | |
| cyclinG1_qrev | CACACAAACCAGGTCTCCAG | |
| tp53_qfor | CCCATCCTCACAATCATCACT | |
| tp53_qrev | CACGCACCTCAAAAGACCTC | |
| mdm2_qfor | GATGCAGGTGCAGATAAAGATG | |
| mdm2_qrev | CCTTGCTCATGATATATTTCCCTAA | |
| puma_qfor | GAACACACGGGTTACAAAGGAC | |
| puma_qrev | GAAAAATCCCAGAGTCTGTAAGTG | |
| elfa_qfor | CTTCTCAGGCTGACTGTGC | |

| | | |
|---|---|---|
| elfa_qrev | CCGCTAGCATTACCCTCC | |
| bactin_qfor | CGAGCTGTCTTCCCATCCA | |
| bactin_qrev | TCACCAACGTAGCTGTCTTTCTG | |
| R217H_SA_F | AAATTGCCAGAGTATGTGTCTGTCC | MspI |
| R217H_SA_R | ATGAGAGCAGCATCATGAAGCAT | MspI |
| R242H_SA_F | ATACCAATCAATTGGACTCATCTCG | ApaLI |
| R242H_SA_R | TTTTCAGCAGTGTAGCAGCAAA | ApaLI |
| p53-R242_sgRNA-2_sense | GTAATACGACTCACTATAGGCTGGGCCGGAGGTCTTTTGGTTTTAGAGCTAGAAATAGC | |
| Rev-sgRNA-scaffold | GGATCCGCACCGACTCGGTGCCACTTTTTCAAGTTGATAACGGACTAGCCTTATTTTAACTTGCTATTTCTAGCTCTAAAAC | |
| R242H-S-Asymm_oligo | C*T*CAATTGTTACATCCTGCTTTTTTGTCCCACAGAGGTCAGTTGCTGGGCCGGAGGTCTTTTGAAGTGCACGTGTGTGCATGTCCAGG*C*A | |
| R242H-KI_for | GGAGGTCTTTTGAAGTGCAC | |
| R242-WT_for | GGAGGTCTTTTGAGGTGCGT | |
| mdm2_MO_F1 | GAGGATGCAGGTGCAGATAAA | |
| mdm2_MO_R1 | CTGCTGTTGTGAGGTAGATGAG | |
| mdm2_MO_F2 | GCAAGGTTGACAACGAGAAAC | |
| mdm2_MO_R2 | CTGTCCGACTTATGCCTCTTC | |
