## Supplementary material for "*tp53* R217H and R242H Mutant Zebrafish Exhibit Dysfunctional p53 Hallmarks and Recapitulate Li-Fraumeni Syndrome Phenotypes": Key Resource Table

**Key resources table**

| REAGENT or RESOURCE | SOURCE | IDENTIFIER |
| --- | --- | --- |
| Antibodies | | |
| Mouse anti-p53-9.1 antibody | Abcam | Cat# ab77813; RRID:AB_10864112 |
| Goat anti-Mouse IgG (H+L) Cross-Adsorbed Secondary Antibody, HRP | Thermo Fisher Scientific | Cat# G-21040; RRID:AB_2536527 |
| Rabbit IgG, anti-phospho-Histone H3 (Ser10) | Millipore Sigma | Cat# 06-570; RRID:AB_310177 |
| Goat anti-Rabbit IgG (H+L) Cross-Adsorbed Secondary Antibody, Alexa Fluor™ 488 | Thermo Fisher Scientific | Cat# A-11008; RRID:AB_143165 |
| Chemicals, peptides, and recombinant proteins | | |
| MEGAshortscript T7 kit | Thermo Fisher Scientific | Cat# AM1354 |
| T7 Endonuclease I | New England Biolabs | Cat# M0302S |
| Camptothecin | Sigma | Cat# C9911 |
| Acridine Orange | Thermo Fisher Scientific | Cat# A1301 |
| Stain-free acrylamide gel | Bio-Rad | Cat# 1610183 |
| Immun-Blot® Low Fluorescence PVDF membrane | Bio-Rad | Cat# 1620261 |
| SuperSignal™ West Dura Ultimate Sensitivity Substrate | Thermo Fisher Scientific | Cat# 37071 |
| Precision Plus Protein™ All Blue Prestained Protein Standards | Bio-Rad | Cat# 1610373 |
| TRIzol reagent | Life Technologies | Cat# 15596026 |
| Direct-zol RNA MicroPrep kit | Zymo Research Corporation | Cat# R2060 |
| Paraformaldehyde | Sigma | Cat# P6148 |
| Bovine serine albumin | Fisher | Cat# SH3057402 |
| Sheep Serum | Sigma | Cat# S22 |
| Collagenase | Sigma | Cat# C0130 |
| FxCycle™ Violet Ready Flow™ Reagent | Thermo Fisher Scientific | Cat #R37166 |
| 10% neutral buffered formalin | Sigma | Cat# HT501128 |
| Paraplast Surgipath paraffin | Leica | Cat# 39601006 |
| Deposited data | | |
| RNA sequencing data | This study | GSE267760 |
| Whole genome bi-sulfite sequencing data | This study | GSE268283 |
| Experimental models: Organisms/strains | | |
| Zebrafish: *casper* (AB) wildtype | University of Ottawa Animal Care and Veterinary Staff | ZDB-GENO-080326-11 |
| Zebrafish: *tp53* null (CG1) | Ignatius et al (2018)^13^ | ZDB-ALT-190529-2 |
| Zebrafish: *tp53* R217H/R217H | Prykhozhij et al (2018)^11^ | ZDB-ALT-190131-7 |
| Zebrafish *tp53* R242H/R242H | This study | N/A |
| Oligonucleotides | | |
| Primers for genotyping mutant lines, amplifying cDNA, and qPCR | This study (Table S2) | N/A |
| CRISPR sgRNAs | This study (Table S2) and Prykhozhij et al (2018)^11^ | N/A |
| Morpholino: MO2-mdm2  5' - TGTTAAGAGATTCAGTACGCACCGC - 3' | GeneTools | ZDB-MRPHLNO-100429-9; described in Parant et al (2010)^20^ |
| Recombinant DNA | | |
| Plasmid: pT3TS-nCas9n | Jao et al (2013)^53^ | RRID:Addgene_46757 |
| Software and algorithms | | |
| R and R studio | Shindelin et al (2012)^56^ | https://www.Rstudio.com |
| Fiji | Github | <https://imagej.net/Fiji>; RRID:SCR_002285 |
| Zen Microscopy Software | ZEISS | RRID:SCR_013672 |
